# Automatic orofacial behaviors remap within cortical substrates for voluntary motor control in freely moving primates

**DOI:** 10.64898/2026.11.06.764525

**Authors:** Elena Ferretti, Francesca Ferrara, Ebrahim Ismaiel, Francesca Lanzarini, Elisabetta Palagi, Monica Maranesi, Luca Bonini

**Affiliations:** Department of Medicine and Surgery – Neuroscience Unit, University of Parma, Parma - Italy; Ernst Strüngmann Institute (ESI) of the Max Planck Society, Frankfurt am Main, Germany; Department of Biology, University of Pisa, Pisa - Italy

## Abstract

Orofacial behaviors, like yawning and emotional facial expressions, involve phylogenetically conserved subcortical circuits. Yet, their context-dependent modulation in primates suggests overlooked cortical contributions. Here, we wirelessly recorded from 640 ventral motor and premotor sites in two freely moving macaques during spontaneous orofacial actions combining automatic and voluntary components to varying degrees (yawning, threat, bite, drink). We identified coupling between synaptic input (local field potential) and spiking output in 169 spatially distributed sites during yawning, while threat display was sparsely represented. Yawn-related sites exhibited an early beta-band suppression followed by high-gamma power enhancement. Of 401 isolated single neurons, 95—predominantly putative medium-sized pyramidal cells—encoded yawning, mostly together with voluntary biting and drinking actions, while showing no modulation during the observation of conspecifics’ yawn. Yawn-related units either switched or maintained their mouth opening/closing tuning across yawning and more voluntary actions, suggesting counteracting and synergistic downstream influences on yawning motor control. In contrast, threat-related units predominantly preserved their opening/closing tuning during biting and drinking. Head-free intracortical microstimulation of sites hosting yawn-related units preferentially evoked localized mouth and face movements, supporting their causal role in orofacial motor control. Finally, projecting neural activity into a low-dimensional manifold showed that voluntarily controlled and largely automatic behaviors engaged distinct population dynamics while preserving the discriminability of individual orofacial displays. These findings reveal that cortical substrates of voluntary motor control can be recruited to shape otherwise automatic orofacial displays.

## Introduction

A variety of orofacial actions, such as yawning and threat displays, largely rely on phylogenetically conserved subcortical circuits (1, 2). Yet their expression can be flexibly modified according to current goals and contextual demands, likely via top-down cortical influences.

Although automaticity and voluntary control constitute coexisting components of most adaptive behavior, classical observations in neurological patients suggest that voluntary and automatic orofacial displays rely on dissociable neural pathways (3): voluntarily generated facial movements depend on the integrity of corticobulbar pathways, whereas emotionally triggered movements can be preserved through anatomically distinct circuits in both humans (4) and non-human primates (5). More recent neurophysiological evidence, however, has challenged a strict separation between these systems, showing that both emotional and voluntary facial gestures can be decoded from neuronal activity in lateral sensorimotor and anterior cingulate cortices (6), suggesting that automatic and voluntary actions may recruit partially overlapping cortical substrates. However, whether and how overlapping cortical representations enable voluntary signals to modulate motor patterns generated primarily by subcortical circuits remains unknown.

Spontaneous orofacial behaviors of freely moving monkeys comprise evolutionarily conserved, highly stereotyped motor patterns that can be shaped by behavioral and social context, thereby providing a particularly suitable model for addressing these questions. Yawning, for example, is a highly stereotyped orofacial sequence (7, 8), conserved across phylogeny—from fishes to mammals, including non-human primates and humans (9)— that emerges during prenatal life (10, 11) and can persist despite extensive cortical damage (12), consistent with its predominant dependence on subcortical and brainstem circuits (13–15). Yet its expression can be adaptively modified: macaques (16) and geladas (17) alter their yawning displays according to social circumstances, for example by selectively exposing or concealing their teeth or combining yawns with different vocalizations (18), while chimpanzees can incorporate additional actions that reduce mouth aperture (19). In humans, even the probability of contagious yawning is strongly influenced by social context (20). Thus, an automatic motor program can remain accessible to flexible, potentially top-down modulation. Threat displays provide a complementary example. Like yawning, they constitute naturally occurring, evolutionarily relevant orofacial behaviors that can be elicited without explicit motor instruction and engage stereotyped patterns of facial and mouth movements (21). Unlike yawning, however, threat displays are generated in response to salient external events (22) and are more closely linked to emotional and social context (22, 23). Thus, despite their automaticity and predominantly subcortical control, these orofacial displays are distinguished by several ethological and neurobehavioral features along the continuum between mostly automatic and predominantly voluntary actions.

Critically, both yawning and threat recruit many of the same orofacial effectors involved in voluntarily controlled actions, such as biting and drinking. Indeed, although automatic motor components contribute to the rhythmic execution of these actions (e.g. chewing and swallowing), their planning and initiation are largely voluntary and dependent on cortical control (24, 25). The ventral motor and premotor frontal cortices are key nodes in the voluntary initiation and control of such orofacial actions (23, 26–28) and project extensively to subcortical nuclei and brainstem centers controlling their execution and rhythmic unfolding (29, 30). This anatomical and functional organization raises the possibility that automatic behaviors are not represented by dedicated cortical motor circuits but instead dynamically recruit neuronal substrates primarily devoted to voluntary motor control. Such recruitment could allow descending signals to voluntarily facilitate, counteract, or otherwise reshape components of ongoing subcortically generated motor patterns.

Testing this hypothesis has been difficult because neurophysiological studies of primate motor cortex have traditionally relied on constrained, head-restrained conditions that limit the expression of spontaneous, species-typical behaviors. Here, we leveraged a recently developed neurobehavioral platform (31) to record neuronal activity from the ventral motor and premotor cortices of freely moving macaques while they spontaneously produced four orofacial actions—yawning, threat displays, biting and drinking—varying along the continuum from essentially automatic to more voluntarily controlled. We found that automatic orofacial behaviors recruited relatively small and partially overlapping subsets of a broader cortical population primarily engaged during voluntary actions. Critically, this recruitment was behavior-specific: during yawning, cortical units displayed temporally structured activity consistent with both counteracting and synergistic influences on different phases of the subcortically generated motor pattern and were located in cortical sites where intracortical microstimulation triggered orofacial movements. In contrast, threat display recruited fewer units that preserved their motor tuning during biting and drinking actions. At the population level, voluntarily controlled and largely automatic behaviors engaged distinct dynamical regimes while preserving the discriminability of individual actions. Together, these findings reveal a cortical mechanism through which automatic orofacial components of behavior are remapped onto cortical substrates for voluntary motor control, providing a framework for understanding how evolutionarily conserved motor programs become flexibly adapted to current behavioral demands.

## Results

### Cortical input-output dynamics during automatic and voluntary orofacial actions

We chronically recorded extracellular activity from 640 sites spanning the ventral primary motor (13,3%) and premotor (86,7%) cortices of both hemispheres in two adult male macaques (Fig. S1). Half of the sites (n = 320) were recorded only once, whereas the remaining 320 were sampled across three or four distinct sessions (Table S1), for a total of 12 sessions (n = 7 in Mk1; n = 5 in Mk2), typically separated by several days. We ensured that resampled channels did not contain the same single neurons recorded in previous sessions, following a recently described methodology (31, 32).

During each session, we recorded neural activity while monkeys moved freely either inside an experimental room specifically designed for neuroethological recordings (the NeuroEthoRoom, NER) (31) or in their home-cage. Several occurrences of voluntary orofacial actions (i.e. drinking and biting) were elicited in each session by offering food or juice to the monkey, and all selected sessions included at least 7 occurrences of spontaneous yawning (Fig. 1A and Table S2). Threat displays were elicited in 9 of the 12 sessions by an unfamiliar experimenter who approached and stared at the monkey before suddenly opening the mouth (23). Synchronized neural and video data were analyzed offline to extract LFP, MUA, and single-unit (SU) activity, aligned with point events corresponding to the behaviors of interest identified using ethological coding software (33) (see Materials and Methods and Table S2).

**Fig. 1.**
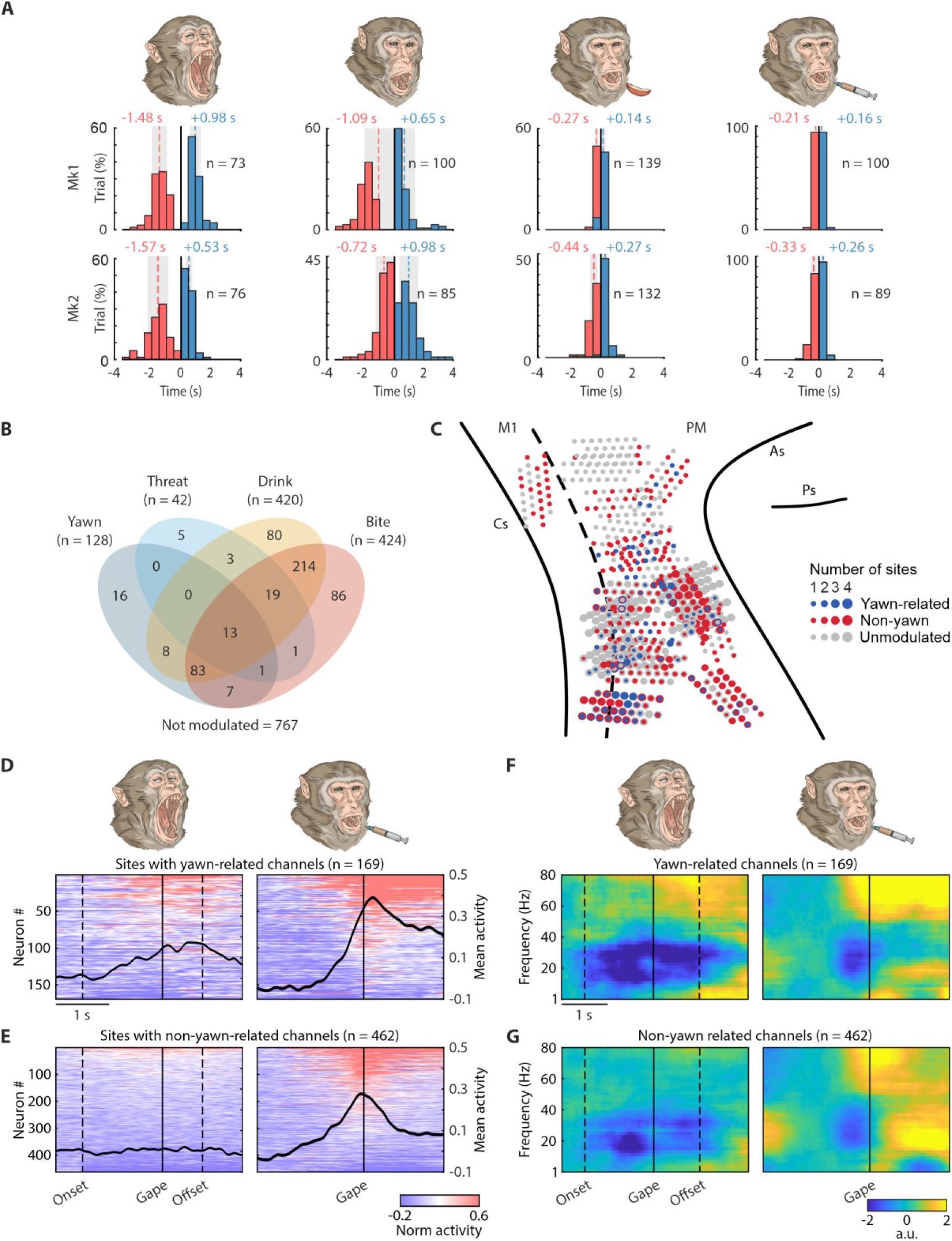
Cortical recording sites, examined behaviors, and associated MUA and LFP modulations. (A) Onset time and duration of mouth opening (negative values, in red) and closure (positive values, in blue) relative to maximal mouth aperture (vertical black line) during yawning, drinking, biting, and threatening. (B) Venn diagram showing the overlap of neurons modulated during yawn, threat, drink, and bite. Numbers indicate the number of neurons within each category and their intersections. Neurons not significantly modulated by any of the behaviors of interest are indicated below (Not modulated, n = 767). (C) Reconstruction of the anatomical location and frequency distribution of channels modulated during yawning (yawn-related) or during orofacial actions other than yawning (non-yawn-related) reported on a template brain obtained by warping the recording sites from Mk1 and Mk2 (see Fig. S1 and Table S1). As, arcuate sulci; Cs, central sulcus; Ps, principal sulcus; M1, primary motor cortex; PM, premotor cortex. (D-E) Heatmaps and corresponding average time courses (black polylines) of normalized activity for yawn-related (D) and non-yawn-related (E) units during yawning (left) and drinking (right) (see also Fig. S2). Vertical dashed lines indicate the average onset and offset of yawn. (F-G) Time course of LFP power modulation across the 1-80 Hz frequency range, averaged across sites with yawn-related (F) and non-yawn-related (G) units during yawning and drinking. Conventions as in D-E.

We isolated a total of 1873 units (1472 MUAs and 401 SUs). Of these, 1303 units (1088 MUAs and 215 SUs) could be tested across a sufficient number of occurrences of all four actions of interest. As a first step, we analyzed their potential modulations during each action. Because no reliable baseline can be identified in freely moving recordings, we adopted a recently proposed approach (31), focusing on 3-second windows centered on the alignment point selected for each action and binning spikes into non-overlapping 250-ms epochs. We then applied a one-way repeated-measures ANOVA (p < 0.01) to test the null hypothesis that each unit’s mean firing rate did not differ across the 250-ms epochs throughout action unfolding (see Materials and Methods). This analysis revealed that 536 of 1303 units (41.1%) were modulated during at least one of the four actions (Fig. 1B, Fig. S2A), with a marked prevalence of selectivity—and, most often, mixed selectivity—for voluntary actions (i.e. drink and bite). Automatic actions showed additional overlap with voluntary-action representations, particularly yawning, which was encoded by 128 of the 536 modulated units. Surprisingly, however, despite the prominent engagement of facial musculature, ethological relevance, and associated arousal of threat displays, only 5 units were selective for threat, and significantly fewer units were modulated during threat than during yawning (n = 42, 3.2%; χ^2^ = 43.51, p = 4.23 × 10^−11^). Furthermore, most threat-related units (28/42, 67%) did not respond during yawning (Fig. S2A; example Unit 1, Fig. S2B), although they showed prominent mixed selectivity for voluntary actions (example Unit 2, Fig. S2B), suggesting substantial segregation between the neural representations of the two automatic behaviors.

Thus, we primarily focused our subsequent analyses on yawning, leveraging the entire dataset. Sites containing yawn-related units (i.e. SUs and/or MUAs; Fig. 1C) were located slightly more frequently in the primary motor than in the premotor cortex (χ^2^ = 4.92, p = 0.03). Interestingly, the same sites responded even more strongly during voluntary ingestive actions, such as drinking (Fig. 1D) or biting (Fig. S3A), showing a peak in population activity corresponding to the transition between the mouth-opening and mouth-closing phases. Another set of 462 sites (31.5%) showed activity related to voluntary orofacial actions but not yawning (i.e. drinking, Fig. 1E, or biting, Fig. S3B), and 835 sites (56.7%) were unrelated to any of the tested actions.

To investigate whether spiking output related to yawning was associated to locally specific synaptic input, we compared the time course of LFP power changes across different frequency bands between yawn-related sites (Fig. 1F) and sites related to orofacial actions other than yawning (Fig. 1G). Yawn-related sites exhibited a marked and early suppression in the beta band (13-28 Hz), followed by a later enhancement in the gamma band (60-80 Hz) during yawning as well as during voluntary mouth actions (see also Fig. S3C, D). This pattern was not evident in non-yawn-related sites, especially during yawning, indicating that the ventral sector of the motor and premotor frontal cortices receives locally specific synaptic input during yawning that, in turn, generates spiking activity associated with both yawning and voluntarily controlled orofacial actions.

Observing others yawn is the strongest known social trigger of yawning (34), suggesting that viewing a yawning individual may recruit cortical mechanisms facilitating or releasing the behavior. Using a previously validated paradigm (35), we tested a total of 780 units (271 SUs and 509 MUAs) while Mk2, seating in a primate chair in head-fixed conditions, was observing videos depicting conspecific yawns or matched control videos showing the same animals immediately before or after yawning (Fig. S4). By applying a 2 x 5 repeated measures ANOVA (factor: Condition and Epoch) to each unit, we found that only the 0.5% of the tested units exhibited a significant, but generally very weak, modulation during yawn observation relative to control stimuli: this proportion was largely below the nominal false-positive rate (α = 0.05), suggesting the absence of modulated spiking activity during yawn observation in this dataset. To further assess whether yawn-related synaptic inputs were present despite being insufficient to generate spiking output, we applied the same analysis to the LFPs recorded from channels exhibiting yawn-related motor activity. No significant modulation was observed in the LFPs (Fig. S4C, D) during the observation of others’ yawns.

Together, these findings reveal that input-output dynamics in the ventral motor and premotor cortex during yawning are specifically related to motor control.

### Yawn-related neurons: putative medium-sized pyramidal cells with mixed selectivity for orofacial actions

To further dissect the cortical mechanisms of yawning as a possible model of voluntarily modulated automatic behavior, we next analyzed single neuron activity. Out of the 401 recorded neurons (Mk1: n = 181; Mk2: n = 220), 95 (23.7%), evenly distributed across premotor (74/328) and primary motor (21/73) cortices (^2^ = 1.27, p = 0.26), were significantly modulated during yawning (Fig. 2A) either selectively (n = 22, 23%; example neuron 1, Fig. 2B) or, more commonly, combined with other orofacial actions (n = 73, 77%; example neurons 2 and 3, Fig. 2B). Yawn-related neurons could peak during the mouth opening (36/95; example neuron 2, Fig. 2B) or closing (59/95; example neuron 3, Fig. 2B) phase of yawning. The remaining neurons were classified as non-yawn-related (n = 165), responding to other tested actions but not yawning, or non-modulated (n = 141), showing no significant modulation for any of the tested actions. Yawn-related neurons with mixed selectivity for voluntary orofacial actions exhibited shorter discharge durations during both drinking (t = 2.6, p = 0.01) and biting (t = 3.9, p = 2.9e^-4^), independent of their discharge duration during yawning (Fig. 2C), suggesting that they temporally rescale their firing to match the dynamics of the encoded orofacial action.

**Fig. 2.**
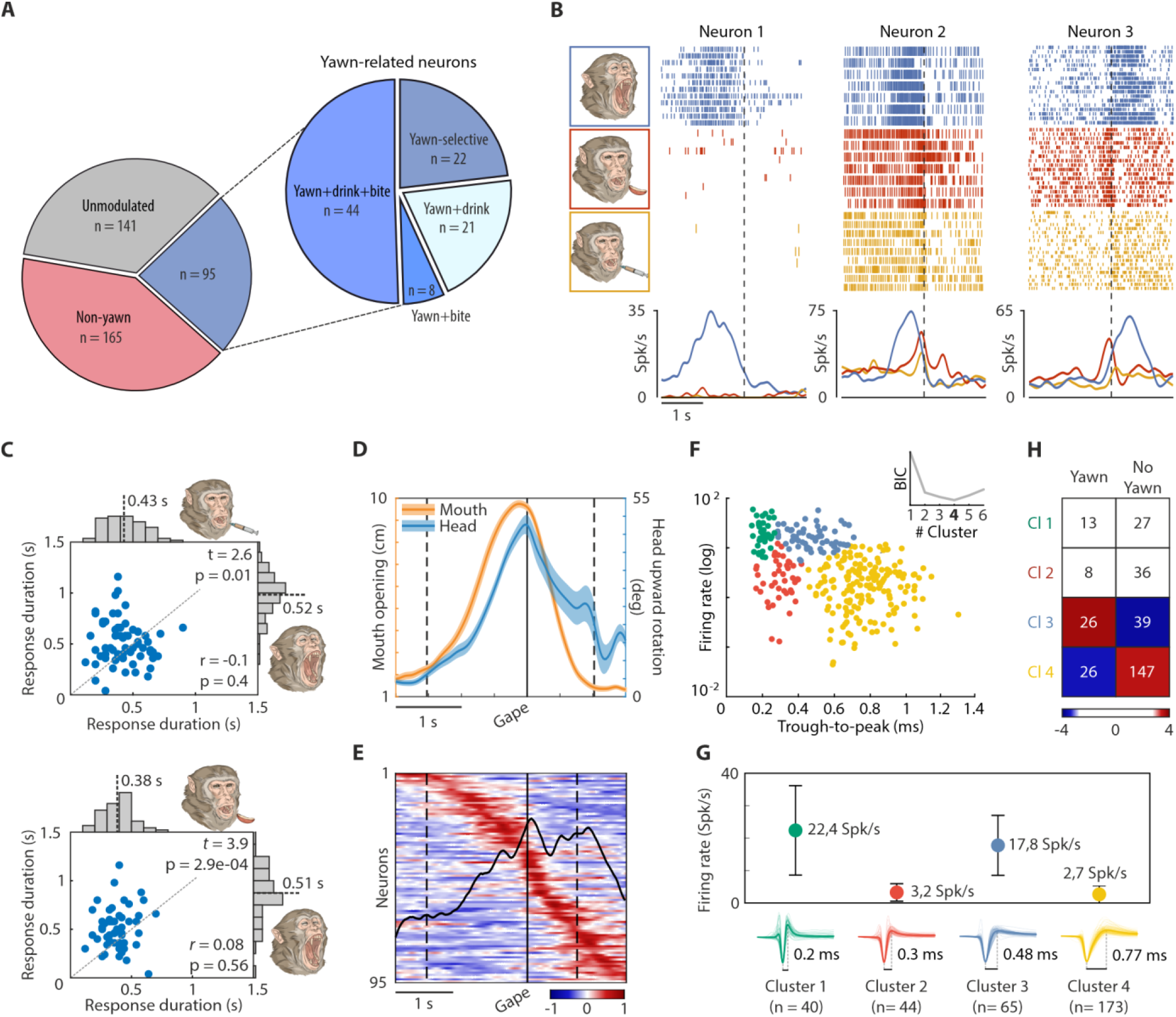
Yawn-related neurons dynamically encode multiple voluntary orofacial actions. (A) Response properties of the recorded neurons. (B) Example neurons selectively modulated by yawning (neuron 1) or jointly by yawning and other orofacial actions (neurons 2 and 3). (C) Response duration of individual neurons during yawning versus drinking (top) and biting (bottom). Dashed line represents the unity line (y = x). Marginal histograms depict the distribution and average discharge durations for each behavior. (D) Average kinematic profiles of mouth opening (orange) and head upward rotation (blue) during yawning (mean ± SE). Dashed lines mark the mean onset and offset of yawning; the black line indicates mouth gape. (E) Heat map showing the net-normalized activity profiles of yawn-related neurons during yawning, from -2 to 1.5 s relative to mouth gape (black vertical line), with the average population response superimposed (black polyline). Neurons are ordered from top to bottom according to the timing of their modulation relative to mouth gape, from the earliest to the latest. Conventions as in D. (F) Two-dimensional distribution of spike waveforms based on trough-to-peak duration and mean firing rate. Colors denote the clusters identified by the GMM, with the optimal number of components (n = 4) determined by the BIC. (G) Waveform shape and firing features of neuronal clusters identified by the GMM. For each cluster, mean firing rate (± SD) and average spike waveform with the corresponding trough-to-peak duration are shown. (H) Heat map of adjusted standardized residuals for yawn-related and unrelated neurons across the four clusters. Color intensity reflects the magnitude of the residuals (−2.73 < residuals < 2.73) while white cells indicate non-significant values (Bonferroni-corrected). Observed frequencies are reported within each cell.

In fact, kinematic measures showed that yawning is particularly long-lasting (mean duration 2.28 ± 0.79 seconds) relative to other orofacial actions, and it is characterized by a wide opening of the mouth accompanied by upward rotation of the head (Fig. 2D). At the population level, yawn-related neurons dynamically encode the whole action unfolding, with 36,8% peaking during the mouth opening phase and the remaining 63,2% during the mouth closure phase. The average peak of population activity corresponded with the maximal mouth aperture (Fig. 2E), thus prompting us to check whether at least part of yawn-related neuron discharge can be explained by somatosensory feedback, which is maximal around the peak of mouth aperture and should vary as a function of movement amplitude. To answer this question we investigated, on a trial-by-trial basis, the possible correlation between individual neurons’ activity and the degree of mouth opening and head upward rotation (Movie S1). The results indicate that yawn-related neurons did not encode nor were specifically influenced by the amplitude of mouth opening or head rotation (Fig. S5), suggesting that they may contribute to higher-level downstream modulation of yawn expression rather than reflecting sensory feedback signals.

Next, we examined the waveform and firing features of yawn-related neurons to investigate whether they could be associated with specific putative cell types (e.g. interneurons vs pyramidal neurons) (36, 37). To this purpose, we clustered neurons using an unsupervised clustering approach (see Materials and Methods) in a two-dimensional feature space defined by trough-to-peak duration and mean firing rate. A Gaussian mixture model (GMM) was fitted to waveform features, and model selection based on the Bayesian Information Criterion (BIC) identified four optimal components, corresponding to clusters (Fig. 2F), ranging from narrow (Cluster 1) to broad (Cluster 4) waveforms with different firing rates (Fig. 3G). Notably, yawn-related neurons were unevenly distributed across clusters (χ^2^=19.61, p=2.05e^-4^), being overrepresented in Cluster 3 and underrepresented in Cluster 4 (Fig. 3H) relative to chance. Based on the established, negative correlation between through-to-peak values and pyramidal cell size (38), these findings suggest that yawn-related neurons mostly correspond to medium-sized pyramidal neurons (39) with particularly high spontaneous firing rate and mixed-selectivity for orofacial actions.

**Fig. 3.**
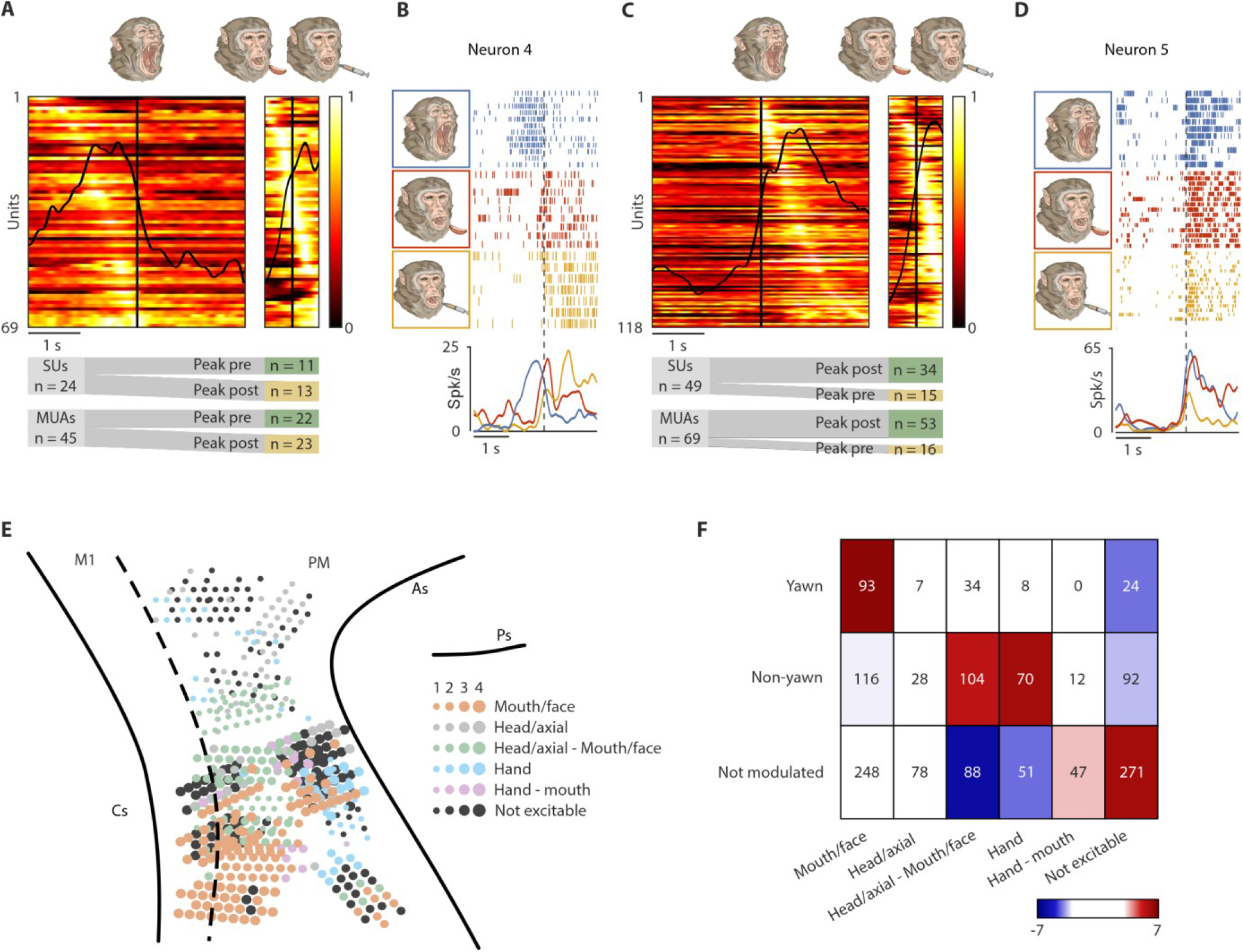
Temporal and causal contribution of yawn-related units to downstream motor control. (A) Net-normalized activity of yawn-related units (SUs and MUAs) peaking during the mouth-opening phase of yawning (n = 69) aligned to the maximal mouth aperture (black vertical line) during yawning (left; -2 to 2 s) and voluntary mouth actions (right; -0.5 0.5 s). In each heat map, units are ordered according to the timing of their peak activity (earliest on top). The alluvial plots below the heatmaps illustrate the proportion of yawn-related SUs and MUAs that either maintained (green) their peak of activity during the mouth-opening phase or switched it (yellow) to the mouth-closing phase across yawning and voluntary orofacial actions. (B) Yawn-related neuron example switching its peak of activity from the mouth-opening phase of yawning to the mouth-closing phase of a voluntary orofacial action (i.e. drinking). (C) Net-normalized activity of yawn-related units (SUs and MUAs) peaking during the mouth-closing phase of yawning (n = 118). Conventions as in (A). (D) Yawn-related neuron example maintaining its peak of activity during the mouth-closing phase of both yawning and a voluntary orofacial action (i.e. biting). (E) Anatomical distribution of ICMS evoked movements (color code) and number of modulated channels in each site (conventions as in Fig. 1A). Head-free ICMS map is obtained using stimulation trains of 500 ms at 100 μA for all but one arrays (in one array in the right hemisphere of Mk2 ICMS was applied with trains of 50 or 500 ms at 10-150 μA. (F) Heat map of adjusted standardized residuals from the significant χ^2^ test comparing the distribution of ICMS evoked movements across sites hosting yawn-related, non–yawn and not modulated units. Color intensity reflects the magnitude of the residuals, with white indicating non-significant values after Bonferroni correction for multiple comparisons (−2.99 < residuals < 2.99). Observed frequencies are reported within each cell.

### Counteracting and synergistic cortical modulation of downstream motor output

Based on the findings reported so far, we hypothesized that the recruitment of cortical neurons associated with the voluntary motor control of orofacial actions during an automatic behavior may counteract or synergistically modulate temporally specific components of the subcortically generated yawn motor pattern. To test this hypothesis, we compared the temporal profiles of all mixed-selective yawn-related units (SUs and MUAs, together providing the most comprehensive index of cortical motor output), across yawning, drinking and biting.

Most units (56%) peaking during the mouth-opening phase of yawning (24 SUs and 45 MUAs) shifted their peak activity to the mouth-closing phase during drinking or biting (Fig. 3A; see also example Neuron 4, Fig. 3B), whereas the majority of units (71%) peaking during the mouth-closing phase of yawning (49 SUs and 69 MUAs) maintained their peak activity in the same phase during drinking or biting (χ^2^ = 13.64, p < 0.001; Fig. 3C), as exemplified by the response of Neuron 5 (Fig. 3D). Together, these findings support the hypothesis that cortical units discharging during the mouth-opening phase of yawning can contribute to a voluntary mouth-closing signal that counteracts the motor output generated by brainstem circuits controlling yawning; conversely, cortical units discharging during the mouth-closing phase of yawning provide a synergistic voluntary signal consistent with the ongoing phase of yawning.

To provide causal support for the hypothesis that the cortical representation of yawning contributes to orofacial motor control, we investigated the effects of intracortical microstimulation (ICMS; Fig. 3E) and their possible relationship with the local neuronal properties. The results (Fig. 3F) showed that sites characterized by yawn-related activity were particularly excitable and their stimulation preferentially evoked orofacial movements (Bonferroni-corrected p = 6.68e^-10^). In contrast, ICMS at sites modulated during actions other than yawning more frequently evoked upper-limb (Bonferroni-corrected p = 2.29e^-08^) and combined axial-orofacial movements (Bonferroni-corrected p = 1.01e^-06^), and sites showing non-modulated activity were predominantly non-excitable (Bonferroni-corrected p = 2.46e^-08^). These results support a possible causal contribution of the motor cortex to the modulation of yawn displays through counteractive and synergistic influences on downstream motor centers involved in the generation of the automatic orofacial display.

In contrast to yawn-related units, threat-related units revealed a predominantly synergistic pattern of activity during threat and the tested voluntary orofacial actions. In particular, threat-related units that did not encode yawning responded predominantly during the mouth-opening phase and exhibited a suppression during the mouth-closing phase (Fig. S6A, top), whereas threat-related units that also discharged during yawning displayed the opposite response pattern, increasing their firing rate preferentially during the mouth-closing phase during both threat and yawning (Fig. S6A, bottom). In both cases, threat-related units maintained their tuning to mouth opening or mouth closing during threat and the other orofacial actions (Fig. S6B; see also Fig. S6C example Unit 3 and Unit 4), suggesting a prevailing synergistic exploitation of neural substrates of voluntary actions for mapping threat displays.

Together, these findings support the idea that different automatic orofacial behaviors are differentially mapped onto motor and premotor neuronal populations primarily dedicated to controlling specific components of voluntary actions, such as mouth opening and closing.

### Population dynamics encode both automatic and voluntary orofacial actions while preserving discriminability

The widespread, combined tuning of motor and premotor units for automatic and voluntary orofacial actions raises a fundamental question: are there dedicated cortical representations for automatic orofacial actions, or do they emerge uniquely from the recruitment of neuronal substrates and dynamics primarily dedicated to encoding voluntary actions?

To address this issue, we focused on the 9 sessions in which all the four actions could be tested in a sufficient number of behavioral occurrences and examined the geometry and dynamics of population activity associated with biting and drinking, on the one hand, and yawning and threat, on the other.

By projecting neural activity into a low-dimensional latent manifold, yawning and threat occupied small, neighboring regions of space, whereas drink and bite were clearly distinct and evolved along more extended and largely parallel trajectories, remarkably similar along the first two principal components but clearly separated along the third (Fig. 4A; Movie S2). Thus, population activity exhibited distinct dynamical regimes for automatic and voluntary actions. This organization was further supported quantitatively using complementary measures of latent geometry. Across sessions, yawning and threat exhibited the smallest trajectory separation among all behavioral pairs (mean normalized trajectory distance = 0.86, *p* = 0.0039; Fig. 4B and S5A, B). The highest latent subspace overlap was observed between biting and drinking (alignment index = 0.35, p = 0.0195), consistent with their stronger functional representation, biomechanical dynamics, and shared ingestive role, followed by yawning and threat (alignment index = 0.27, p = 0.0273; Fig. S5 A, B). Importantly, similar latent dynamics are evident even when considering individual recording sessions, and leave-one-session-out control analysis demonstrated that both trajectory distances and alignment indices were preserved after removal of individual datasets (Fig. S7C, D).

**Fig. 4.**
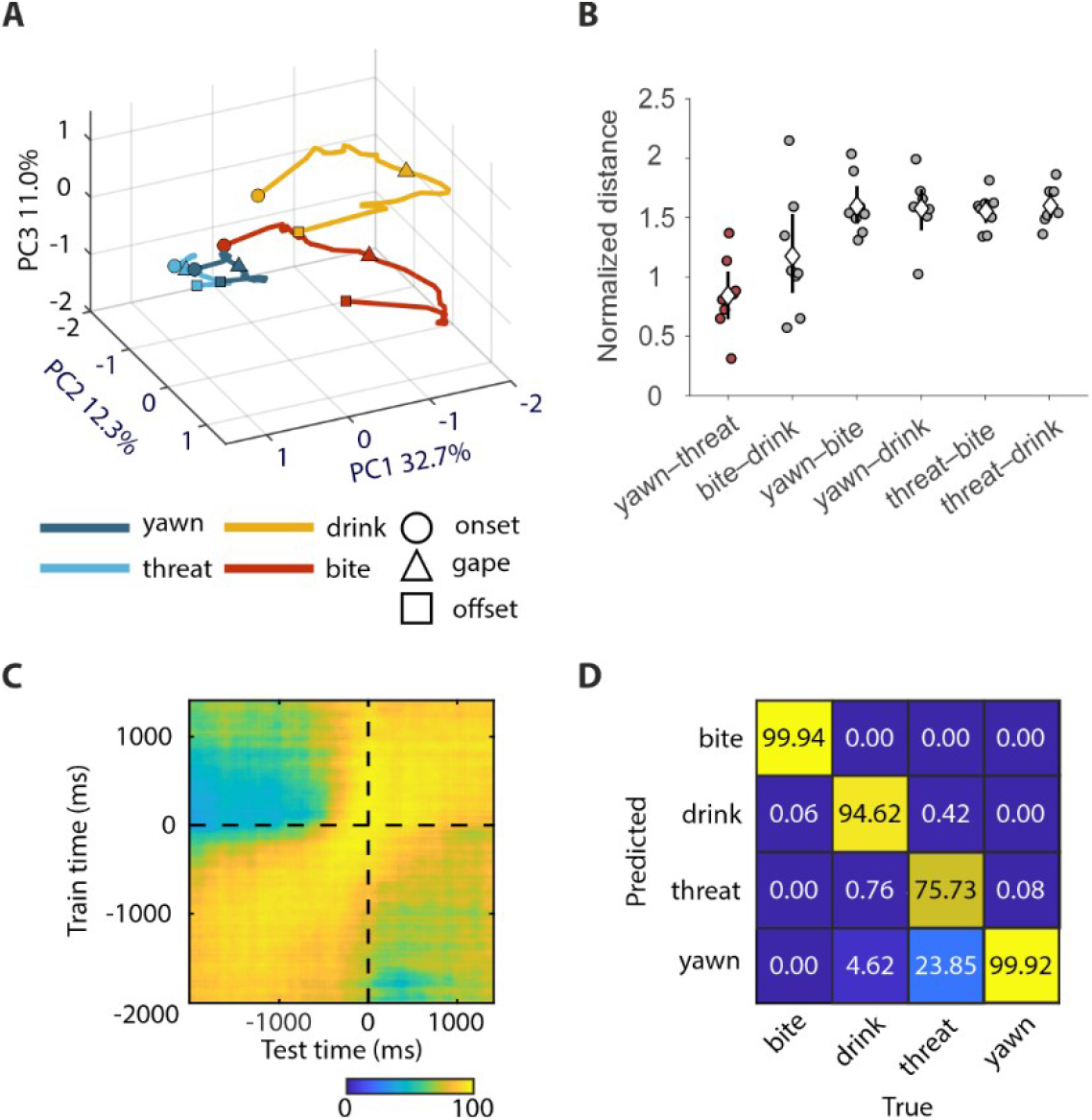
Partial overlap in the neural representation of automatic (threat and yawning) and voluntary (bite and drink) orofacial actions. (A) Pseudo-population latent trajectories from action onset (circles), through gape (triangles) to action end (squares), reconstructed from the activity of all recorded units (SUs and MUAs) pooled across sessions and monkeys during all actions. Population activity was projected onto the first three principal components (PC1–PC3) of the phase-normalized neural manifold (variance explained is indicated along each axis). (B) Normalized trajectory distance for each action pair across recording sessions. Circles represent individual-session values, diamonds indicate across-session means, and vertical bars indicate 95% confidence intervals. Dark red identifies action pairs showing significantly smaller distances in trajectories relative to the other action pairs (one-tailed Wilcoxon signed-rank test, FDR-corrected, *q* < 0.05). (C) Cross-temporal decoding matrix showing classification accuracy for the four actions (yawning, threat, biting, and drinking) using all recorded units pooled across sessions. (D) Averaged decoding results across the analyzed time window (−2000 to +1400 ms). Values represent the average classification performance at matched training and testing times in (C). Columns correspond to true classes and rows to predicted classes; values are column-normalized percentages. Decoding performance was significantly above chance (permutation test, p < 0.001).

Notably, the substantial overlap between yawning and threat, on the one hand, and bite and drink, on the other, did not impair neural discriminability within the two action classes. Indeed, time-resolved cross-temporal decoding analyses using all recorded units (see Materials and Methods) revealed robust (> 75%) and dynamically evolving decoding accuracy for all actions (Fig. 4C), with misclassification mostly reflecting confusion between yawning and threat (Fig. 4D).

Together, these findings reveal that automatic orofacial behaviors remap onto a relatively small, partially shared subset of cortical units primarily dedicated to representing voluntary orofacial actions, reconfiguring population dynamics while preserving both the distinction between automatic and voluntary behaviors and the discriminability of individual actions.

## Discussion

As early as 1837, Heinrich Magnus described a patient who could not open the mouth on command but retained the ability to do so spontaneously during yawning or crying (3, 40). Later lesion studies attributed volitional facial movements to corticobulbar pathways and emotionally evoked ones to separate routes (4) (5), while recent studies in head-restrained monkeys have shown that both emotional and voluntary facial gestures can be decoded from lateral sensorimotor and anterior cingulate activity (6), favoring a distributed cortical network over segregated circuits (41). The present findings help reconcile this discrepancy and shed new light on the mechanisms for the cortical modulation of automatic components of behaviors.

By recording from freely moving monkeys producing spontaneous orofacial displays along the continuum from mostly automatic (i.e. yawning and threat) to predominantly voluntary (i.e. biting and drinking) actions in an ethologically and ecologically relevant context (42), we showed that all four actions can be decoded with similar accuracy from the ventral motor and premotor cortices, confirming recent findings (6) and extending them to yawning and drinking. Nonetheless, voluntary actions engaged substantially larger neuronal populations than automatic actions, and threat displays were significantly less represented relative to yawning, suggesting that cortical recruitment scales with the degree of voluntary motor control rather than movement complexity. Furthermore, we found that yawning, long regarded as a prototypical automatic behavior generated by conserved subcortical circuits (43), is robustly represented at multiple scales by coordinated synaptic and spiking activity, which can hardly be fully accounted for by afferent sensory feedback associated with stretching of the skin and muscles during gaping (7). Sensory feedback is an integral component of motor control, even at the level of central pattern generators, where proprioceptive inputs contribute to shaping and adjusting motor output (44). Our kinematic findings do not exclude a contribution from afferent inputs, but the absence of a relationship between movement amplitude and neuronal response magnitude allows us to rule out the possibility that the recorded activity is simply a reflection of movement-related somatosensory feedback, favoring the interpretation of yawn-related activity as an efferent motor signal.

Notably, neither spiking activity nor LFPs were modulated during exposure to videos of yawning conspecifics in the tested subject. Although the yawn observation experiment was limited to a single animal, this finding is consistent with the hypothesis that contagious yawning may rely predominantly on subcortical mechanisms, as proposed for mimicry phenomena (45), rather than on cortical networks (46). This possibility is supported by the remarkably slow temporal dynamic of behavioral contagion (47) and recent demonstrations of contagious yawning in fishes (48), although the underlying neural circuitry and its evolutionary history remain unknown.

Yawning is thought to be triggered by a subcortical network linking the central nucleus of the amygdala (49), the bed nucleus of the stria terminalis (50), and the hypothalamic paraventricular nucleus, the latter representing the main site for yawning induction (43, 51). However, none of these structures receives direct projections from the ventral motor and premotor cortices. Instead, these cortices project extensively to the reticular formation and the facial and trigeminal nuclei (52, 53), which control the large set of muscles involved in both yawning and voluntary orofacial actions (54). Although several body parts and muscle groups can jointly contribute to yawns and orofacial actions, most yawn-related cortical neurons are putative pyramidal cells primarily tuned to voluntary orofacial actions and located at sites where ICMS evokes orofacial movements, suggesting a causal role in modulating brainstem motor centers specifically related to orofacial motor control. Our finding that a substantial fraction of yawn-related neurons peaking during mouth opening shifted their peak firing to the mouth-closing phase during voluntary actions suggests that descending voluntary signals associated with mouth closure can counteract brainstem motor circuits during the initial phase of yawning. In contrast, yawn-related neurons peaking during mouth closure consistently peaked during the mouth-closing phase of voluntary orofacial actions, thereby providing a synergistic signal to brainstem motor circuits. Although the relationship between neuronal firing and mouth opening or closing across behaviors may reflect differences in biomechanical and postural demands rather than a fixed motor effect, and therefore does not uniquely establish counteracting or synergistic mechanisms, these findings suggest a potential role for the premotor cortex in sculpting an otherwise automatic behavior by recruiting neural resources dedicated to voluntary motor control.

Why is threat, an emotionally triggered and equally automatic response, much less strongly represented than yawning at the cortical level? Our findings show that, like yawning, threat is remapped onto neural substrates primarily dedicated to encoding voluntary orofacial actions and occupies a position in neural space that is close to yawning yet largely segregated from voluntary actions. However, threat-related units largely maintain the same mouth opening/closing tuning across threat and the other tested actions, suggesting that threat displays are not counteracted by direct cortical top-down signals. This is likely because the time course of threat displays is too rapid to be sculpted by cortical activity, making them subject to stronger anticipatory inhibitory influences mediated by prefrontal–amygdala pathways (55, 56).

The present findings support a framework in which the cortical representation of orofacial actions is widely distributed, while still adhering to precise organizational principles. Evolutionarily ancient, automatic motor patterns that rely on subcortical networks can be flexibly regulated through the recruitment and dynamic reconfiguration of upstream motor centers, as previously demonstrated for vocal control across mammalian species (57, 58). This model is consistent with neurological evidence showing that denervation of brainstem centers can lead to pathological yawning, supporting the existence of descending cortical inhibitory influences (15). Such an architecture allows motor signals generated by cortical regions primarily involved in voluntary motor control to facilitate or suppress the output of brainstem motor circuits responsible for automatic behaviors, thereby making them flexibly adaptable to the current context.

## Materials and Methods

### Animal Subjects

Two male rhesus macaques (*Macaca mulatta)*, Mk1 (8 years old, 12 kg) and Mk2 (10 years old, 13 kg) were used in this study. The animals were housed in facilities providing cage space exceeding the requirements of Italian and European regulations and had daily access to an enriched environment including wooden structures and a variety of toys. All experimental procedures complied with the European Directive 2010/63/EU on the protection of animals used for scientific purposes, were approved by the Veterinarian Animal Care and Use Committee of the University of Parma and were authorized by the Italian Ministry of Health.

### Surgical procedure

Each monkey underwent three surgical procedures performed under aseptic conditions. General anesthesia was induced with ketamine (4.5 mg/kg) and medetomidine hydrochloride (0.05 mg/kg), maintained with inhaled isoflurane (IsoFlo, 100% p/p), and physiological parameters were continuously monitored using a multiparameter monitor. Following each procedure, the animals received postoperative antibiotic, anti-inflammatory, and analgesic therapy.

The fMRI-guided stereotactic surgeries involved implantation of a titanium head-post and, subsequently, of a variable number of 32-channel floating microelectrode arrays (FMAs; Microprobes for Life Science, Gaithersburg, MD, USA) in the left and right hemispheres during separate surgeries. The specific insertion sites for the FMAs were determined based on visible anatomical landmarks, particularly the superior and inferior arcuate sulci and the central sulcus. The left hemisphere was implanted first in each animal (n = 4 arrays in Mk1; n = 6 arrays in Mk2, of which 4 were used for this study); next, we implanted the right hemisphere (n = 6 arrays in both monkeys). The implants targeted the ventral part of the lateral precentral cortex (Fig. S1).

After array implantation and bone flap repositioning, a customized recording chamber with a regularly spaced grid was secured to the skull using bone screws and dental cement. The chamber housed the 32-channel Omnetics connectors and the cables from the implanted arrays. During recording sessions, an additional protective cover was attached to accommodate and protect the neural data logger and battery pack, as previously illustrated elsewhere (31).

### Apparatus and behavioral paradigm: freely moving recordings

To record spontaneous behaviors under naturalistic, freely moving conditions, we collected neurobehavioral data across two experimental contexts: the NeuroEthoRoom (NER) and the monkeys’ home cages.

The NER is a transparent Plexiglass enclosure specifically designed for neuroethological recordings (W x H x D, 205 x 205 x 180 cm) and described in detail elsewhere (31). Animals were continuously monitored using eight synchronized high-resolution cameras positioned around the enclosure and recording at a sampling rate of 50 Hz. The environment was enriched with a variety of items to promote spontaneous species-specific behaviors. Recording sessions in the NER lasted an average of 40 minutes.

In the home-cage (W x H x D, 153 x 144 x 253 cm), video recordings were acquired at a 50 Hz using two cameras positioned at different angles in front of the cage, providing full coverage of the cage volume. Home-cage recording sessions lasted between 2 and 9 hours and included some overnight recordings. During overnight sessions, infrared cameras were used together with three infrared light sources positioned approximately 2 m from the cameras to ensure adequate illumination while remaining invisible to the monkey.

Each monkey was tested individually. At the beginning of each recording session, the animal, previously trained to do so, entered the primate chair directly from its home cage to allow connection of the data logger. Four reflective markers were then attached to the headpost to track the monkey’s head movements during free behavior. Subsequently, the monkey was either released back into the home cage or transported to the laboratory and released into the NER.

In both settings, the experimenter interacted with the monkey to elicit voluntary orofacial actions and automatic orofacial behaviors. To elicit biting, the animal was offered small food morsels either directly into its mouth, by tossing them onto the floor, or by suspending them from the ceiling using transparent lines. Juice or water was delivered via a plastic syringe to induce drinking. To elicit threat behaviors, an unfamiliar experimenter entered the room and stared at the monkey, occasionally opening the mouth, which typically induces an open-mouth threat display in the animal. Following these interactions, the monkey was allowed to move freely and explore the environment. Only sessions in which the monkey displayed at least seven spontaneous yawns were included in the dataset, resulting in a total of 12 recording sessions (Mk1: 7 recording sessions in the NER; Mk2: 5 recording sessions in the home-cage).

### Apparatus and behavioral paradigm: head-fixed recordings during yawn observation

To investigate whether neuronal activity was modulated by the observation of yawning in conspecifics, the monkey was head-fixed while viewing video stimuli presented on a 32-inch monitor positioned 70 cm away (Fig. S4). Each recording session consisted of two experimental blocks: one *yawn video experimental block* and one *control video experimental block*. Each experimental block followed the same structure, consisting of a 3-minute black screen, a 5-minute video presentation, and a 3-minute black screen. The order of presentation of the two experimental blocks was randomized across sessions.

During the 5-minutes video presentation periods, the monkey viewed video clips depicting either familiar conspecifics (i.e., macaques housed in the same facility) or unfamiliar conspecifics. The unfamiliar stimulus set comprised 21 clips (5 s each), each presented twice, whereas the familiar stimulus set comprised 20 clips (3 s each), each presented three times. Clips were separated by a variable (1-3 s) black-screen inter-trial interval, resulting in a total video of 5 min.

In the yawn block, clip depicted conspecifics yawning, with yawning beginning 1 s after clip onset. In the control block, each clip was paired with its corresponding yawn clip and depicted the same macaque in the same setting for the same duration while engaged in non-yawning behavior, filmed immediately before or after the corresponding yawn episode, thereby ensuring that subject identity, background, and other visual features were matched across conditions. Thus, the only difference between paired clips was the presence or absence of yawning.

During each recording session, the monkey’s behavior was continuously recorded by two cameras, one positioned immediately above the stimulus monitor to provide a frontal view of the animal. The recordings were subsequently inspected offline using BORIS. Periods in which the monkey moved, closed its eyes, or clearly shifted its gaze upward or laterally away from the monitor were annotated as state events. Trials overlapping with these periods were excluded from the neural analyses, retaining only those in which the monkey remained still, with its eyes open and its gaze directed toward the stimulus monitor. Given the known visual response properties of ventral premotor neurons during action observation (59), these conditions should minimize the influence of differences in over visual attention on the results.

### Recording technique

We simultaneously recorded from 128 channels distributed across four floating microelectrode arrays (FMAs) in all but one session (in which we recorded from 64 channels), using a wireless neural data logging system (RatLog-128, Deuteron Technologies, Jerusalem, Israel). The recordings were performed with a bandpass filter set to 1– 7000 Hz and a sampling rate of 32000 Hz per channel. Neural signals were amplified, digitized, and stored locally on a MicroSD memory card. The logger was powered by an external 3.7 V battery directly connected to it and housed within a protective cover (31).

Neural activity and video recordings were synchronized in both the NER and the home cage using hardware-generated trigger signals. In the NER, a transceiver delivered a continuous 5 V, 50-Hz synchronization pulse generated by an external PC equipped with a National Instruments data acquisition board. The pulse was recorded by the neural logger together with the neural data and simultaneously fed to the SIMI Motion capture system (Simi Reality Motion Systems GmbH, Unterschleißheim, Germany), where it triggered frame acquisition from the eight synchronized cameras. In the home cage, a continuous 5 V, 50-Hz square pulse was generated by MonkeyLogic (60) and delivered through a movable National Instruments data acquisition board to two Basler video recording systems (Basler AG, Ahrensburg, Germany), each controlling one camera. A corresponding digital synchronization signal was simultaneously recorded by the neural logger together with the neural data. In both settings, the synchronization signals enabled precise offline alignment of neural activity and video recordings.

The same recording setup and synchronization procedure used during home-cage recording sessions were also employed during the head-fixed observation task. MonkeyLogic was additionally used to present the yawn and control videos on the stimulus monitor. At the onset and offset of each stimulus block, a TTL pulse generated by MonkeyLogic and delivered through the National Instruments data acquisition board was recorded by the neural logger together with the neural data, enabling precise offline alignment of neuronal activity with stimulus presentation.

### Mouth and head kinematics extraction during yawn

Mouth and head kinematics during yawning were extracted from video recordings using either the video-based motion tracking implemented in SIMI Motion (SIMI Reality Motion Systems, Germany) or the markerless pose estimation toolbox DeepLabCut (61). Upper and lower lips were tracked to quantify mouth kinematics, whereas markers mounted on the headpost (Fig. S5A) were used to reconstruct head movements (see Movie S1). In SIMI-based sessions, landmarks were manually annotated every 100 ms and linearly interpolated to obtain trajectories sampled at 20 ms resolution. In DeepLabCut-based sessions, the same landmarks and headpost markers were automatically tracked using a trained deep neural network. Trials in which the lips or headpost markers could not be reliably tracked at 100 ms intervals because of temporary occlusions or suboptimal camera views were excluded from analysis.

For each trial, mouth opening was quantified as the Euclidean distance between upper and lower lips across time (Fig. S5C). Head elevation was quantified as the angular displacement of the head within the sagittal plane relative to its initial orientation. In particular, at each frame, head direction vector was estimated from the midpoint between the left temporal marker (*L*) and the right temporal marker (*R*), and a third anterior/posterior head marker (*A*) (located either anteriorly or posteriorly, depending on marker visibility within each trial).

To quantify yawning-related head elevation while minimizing contamination from lateral head movements, for each frame *t*, the midpoint between the bilateral temporal markers was computed as:

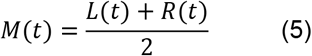

A raw head axis vector, *H*(*t*), was defined between the temporal midpoint and the third head marker as

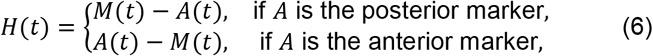

and normalized to obtain the unit head axis,

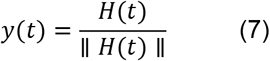

At the first valid frame *t*_0_, a baseline head reference frame was constructed. The normalized left–right axis was defined as

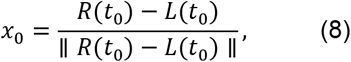

the baseline head axis as

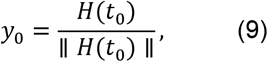

and the third orthogonal axis as

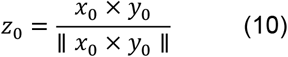

The vectors *y*_0_ and *z*_0_ define the baseline sagittal plane, whereas *x*_0_ represents the orthogonal left–right direction. To isolate upward–downward head motion, the current head axis was projected onto the baseline sagittal plane by removing its component parallel to the baseline lateral axis *x*_0_,

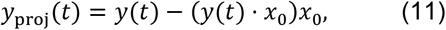

and normalized,

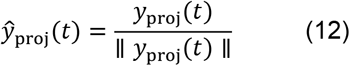

The instantaneous head tilt angle was then computed within the baseline sagittal plane as

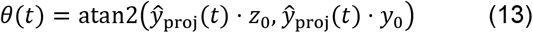

Finally, head tilt was expressed relative to the initial posture as

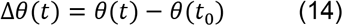

### Spike sorting and assessment of the recording stability

All analyses were performed offline. Signal from all the recorded channels was band-pass filtered (300-6000 Hz), and spikes were detected using a negative threshold of 3 standard deviation from the mean signal of each channel. After possible artefacts removal, detected spikes constituted the multiunit activity (MUA). Detected spikes were then sorted using either the automated algorithm MountainSort (62), setting “noise overlap” threshold at 0.20 for single units, or Plexon Offline Sorter (Plexon Inc). Putative single units (SUs) were subsequently validated by manual curation and by assessing the temporal stability of spike isolation according to previously described criteria (31). In addition, for two pairs of sessions from the left hemisphere of Mk1 collected 7 days apart, we verified that the same neuron was not included more than once across sessions. When duplicate recordings were identified, only the unit with the highest signal-to-noise ratio was included in the final dataset.

### Freely moving behavior and single unit analysis

Behavioral data were analyzed at a 50 Hz frame-by-frame resolution using BORIS (63). Two expert observers, with extensive experience in non-human primate behavior within laboratory settings, independently performed the scoring. The observations relied on an ethogram developed specifically for both NER and home-cage recording sessions (Table S2). In particular, for each behavior of interest, we identified a key temporal event used as the alignment point, corresponding to the transition between distinct motor phases. For transitive actions, the alignment point corresponds to the moment when a body part makes contact with an external object (e.g., food during biting, or the syringe during drinking). In contrast, for intransitive actions such as yawning or threat displays, the alignment point (i.e., maximal mouth aperture) represents the moment of maximal somatosensory feedback.

For each recording session, to determine whether a unit (single unit and/or multi-unit) was modulated in relation to a specific behavior, we selected a time window of 3 seconds centred on the event alignment point (1.5 s before and 1.5 s after) avoiding the inclusion of other behaviors occurring before and/or after. The spikes of each SU and/or MU within these time windows were binned into non-overlapping epochs of 250ms and a one-way repeated measures ANOVA was conducted across the 250ms time epochs for each behaviour separately, under the null hypothesis that if a unit is not modulated for a certain action, then its activity should remain unchanged during action unfolding.

Because behavioral occurrences vary greatly under freely moving conditions, we wanted to prevent unbalanced statistical power across behaviors. To minimize false positives in behaviors with a greater number of trials, we repeated the described ANOVA 1000 times, each time randomly selecting a number of trials equal to that of the least represented behavior in the session. A unit was considered significantly modulated for a specific behavior if 60% of the ANOVA iterations resulted in a significant main effect (α = 0.01).

### Response duration during yawning and other orofacial actions

To compare response duration during yawning and other orofacial actions, neural activity from yawn-related units was extracted in 20 ms bins within a time window of −2 to 1.5 s relative to mouth maximum opening during yawning, and −1.5 to 1.5 s relative to mouth maximum opening during biting and drinking. For each neuron, response duration in relation to a given action was defined as the interval between the first pre-peak time point at which neural activity exceeded the mean firing rate by 1 standard deviation and the first post-peak time point at which activity dropped below the same threshold. Paired sample t-tests were used to compare response durations between action pairs, while Pearson correlation analyses were used to assess the relationship between neural response durations across behaviors (Fig. 2C).

### Correlation between neuronal responses and yawn kinematics

To assess the relationship between neuronal response magnitude and yawn kinematics, for each neuron the firing rate was computed for every yawn trial within a time window extending from −1.5 to +1.5 s relative to the alignment point (maximum mouth opening). Spike counts were calculated by binning single trials activity (Gaussian filter σ = 3) in 60 ms bins with 20 ms step, to reduce the influence of noise in single-trial firing rate estimates. For each yawn trial, Kendall’s rank correlation was computed between peak firing rate, on one side, and maximum mouth opening (Fig. S5G) and maximum head elevation (Fig. S5H), on the other.

### Temporal dynamics of neuronal responses during behavior

To characterize the temporal dynamics of neuronal responses, heat maps of net-normalized firing activity averaged across trials were generated for single units (SUs) and/or multi-units (MUAs) during the behaviors of interest. These analyses were used both to characterize the temporal dynamics of units responsive to automatic behaviors (i.e., yawning and threat) and to compare their response profiles during these behaviors with those observed during voluntary orofacial actions (i.e., biting and drinking). For each unit, firing rates were computed in 20-ms bins within a behavior-specific time window aligned to the moment of maximum mouth opening. Net normalization was performed by subtracting the mean firing rate within the analyzed window and dividing by the maximum firing rate observed within the same window. When the activity of a unit was examined during voluntary orofacial actions and the unit was responsive to both biting and drinking, only the action associated with the higher mean firing rate was retained for comparison.

### Waveforms clustering analysis

#### Extraction of single units waveforms

The average waveform of single units was computed by randomly selecting 1000 spikes from the unfiltered wideband signal, using a 3 ms window centered on the spike’s absolute minimum (1 ms before and 2 ms after). Each waveform was spline interpolated to obtain 1000 points within the 3 ms window, independent of the original sampling rate, and then realigned to its absolute minimum. This process yielded the final average waveform for each unit. From the final dataset, we excluded units with: 1) noisy non-canonical waveforms (multipeak, e.g., waveforms exhibiting multiple local maxima between the main trough and the subsequent peak) (n = 4); 2) a main trough amplitude smaller than that of the nearest peak, whether occurring before or after the main trough (n = 34), as these units likely correspond to axonal fibers (64) (n = 42) ; and 3) waveforms lacking a post-main-trough peak, which prevented computation of the trough-to-peak duration (n = 33). The final dataset comprised 322 units.

#### Clustering of single unit waveforms

For each single unit, average spike waveforms were smoothed, and the trough-to-peak duration was computed as the temporal interval between the negative deflection of the waveform and the subsequent positive peak. The mean firing rate was calculated as the total number of spikes recorded during the session divided by the overall session duration.

To identify clusters based on these two electrophysiological features, the two-dimensional data points were fitted using a Gaussian mixture model (GMM; MATLAB function: fitgmdist). Model selection was performed using the Bayesian Information Criterion (BIC). GMMs with one to six Gaussian components were fitted to the two-dimensional feature space (trough-to-peak duration and mean firing rate) using both full and diagonal covariance matrices. BIC values were compared across all tested models, and the model with the lowest BIC was selected. This procedure identified a GMM with four components and a diagonal covariance structure as the best-fitting model (Fig. 2F).

### LFP Processing and Time-Frequency Analysis

Using custom software written in MATLAB R2023b (MathWorks, Natick, MA, United States), local field potential (LFP) signals were downsampled to 1 kHz prior to analysis and band-pass filtered between 0.5 and 200 Hz using a zero-phase second-order Butterworth filter, followed by a zero-phase notch filter centered at 50 Hz (quality factor = 35). Filtering was performed bidirectionally to avoid phase distortions. Time-frequency decomposition was computed using the multitaper spectral estimation framework implemented in the Chronux toolbox (http://chronux.org). Spectrograms were generated using a sliding window of 500 ms advanced in 50 ms steps, with discrete prolate spheroidal sequence tapers (time-bandwidth product TW = 3, number of tapers K = 5). Spectral power was estimated over the frequency range of 0.5-200 Hz and subsequently converted to logarithmic units (10·log10 power). To facilitate comparisons, spectrograms were normalized relative to a trial-specific pre-onset period (−1.5 s to −0.5 s preceding movement onset). At each frequency bin, pre-onset mean and standard deviation were computed and used to z-score spectral power across time. To further standardize dynamic range across recordings, normalized values were linearly rescaled independently for each frequency band to the interval (−1, 1). Each trial was segmented according to its annotated event sequence, and the time–frequency representation within each segment was interpolated onto a standardized temporal grid consisting of 100 equally spaced bins per behavioral epoch. The resulting normalized spectrogram preserved the relative progression of behavioral states while minimizing variability arising from differences in absolute movement duration. Segment-wise, normalized representations were subsequently concatenated to obtain a common trial trajectory for population-level analyses.

For visualization, trial-normalized LFP spectrograms were averaged within each condition and channel class and displayed on an event-aligned normalized time axis. Band-limited power was quantified within the 1–8 Hz (low-frequency), 13–28 Hz (beta), and 60–200 Hz (high-gamma) bands. Channel-level modulation indices were computed as the difference between band-limited power during the behavioral epoch of interest and the corresponding pre-onset epoch. During yawn execution, modulation values were averaged across trials within each recording channel. During yawn observation and control observation, modulation values were averaged within each channel–stimulus combination. Statistical significance of modulation relative to pre-onset was assessed using two-sided Wilcoxon signed-rank tests. Differences among conditions were evaluated using Kruskal–Wallis tests, and uncertainty of median modulation estimates was quantified using bootstrap-derived 95% confidence intervals (10,000 resamples). Statistical significance was defined as p < 0.05.

### Latent population dynamics and trajectory analysis

To examine the population geometry underlying the tested actions we analyzed neural activity from all available single- and multiunits in relation to three main events consistently identified across actions: mouth-opening onset, maximal gape, and mouth closing offset. For each unit, firing rates were computed using adaptive phase-normalized windows spanning the interval from mouth-opening onset to maximal gape and from maximal gape to mouth closing offset. The opening and closing phases were each resampled into 25 non-overlapping phase bins, yielding a common phase axis of 51 samples (0 = onset, 0.5 = maximal gape, 1 = offset). Activity was smoothed using a one-dimensional Gaussian kernel (σ = 1 phase bin) and soft-normalized by dividing ach neuron’s firing rate by its absolute maximum across all trials and conditions considered in a session, plus a constant of 5 spikes/s. Population trajectories were first constructed independently within each recording session using only neurons simultaneously recorded in that session. Trial-averaged neural activity matrices from all behaviors (phase samples × neurons) were concatenated to construct a session-specific population matrix. Neuronal activity was mean-centered across samples before principal component analysis (PCA) calculation. Six principal components were initially computed for each session. The first five principal components were used for subspace alignment analyses, whereas the first three principal components were used for trajectory visualization and trajectory-based similarity metrics.

To quantify similarities between behavioral trajectories, two complementary metrics were computed for every pair of actions within each session. The alignment index measured overlap between low-dimensional neural subspaces and was calculated using the covariance structure of activity projected onto the first five principal components. The overlap between the neural subspaces associated with actions *A* and *B* was quantified as:

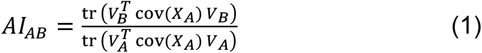

where: *X*_*A*_ and *X*_*B*_ are the phase-normalized population activity matrices for actions *A* and *B*(phase samples × neurons), cov(*X*_*A*_) is the covariance matrix of action *A, V*_*A*_ and *V*_*B*_ are the first five principal-component loading vectors extracted from *X*_*A*_ and *X*_*B*_, respectively, and tr(·) denotes the matrix trace. The alignment index ranges from 0 (orthogonal subspaces) to 1 (identical subspaces).

Normalized trajectory distance quantified the mean Euclidean separation between trajectories after variance normalization. After projection onto the first three principal components, each behavior was represented by a trajectory:

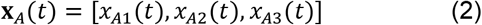

sampled at *T* = 51 normalized phase points. To facilitate comparisons of trajectory geometry across behaviors, latent trajectories within each recording session were normalized by the square root of the total variance of the session-wide latent population activity, where x_k^session denotes the projection of the pooled session-wide latent activity onto principal component k (computed across all behaviors after projection onto the first three principal components):

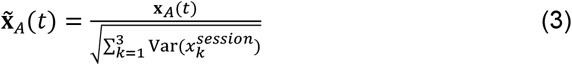

The normalized trajectory distance between actions *A* and *B* was then computed as the mean Euclidean separation across the normalized phase axis:

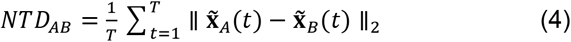

where ∥·∥_2_ denotes Euclidean norm. Smaller values indicate similar latent trajectories, whereas larger values indicate greater separation in neural state space.

To determine whether observed geometries generalized across recording sessions, we performed leave-one-session-out (LOSO) analyses. Session-specific metrics were compared with the corresponding metrics computed from pooled datasets in which the respective session had been excluded, using linear regression and Pearson correlation analyses.

For visualization purposes, pooled pseudo-population trajectories were generated by concatenating phase-normalized units across sessions without assigning or imputing activity for neurons not recorded in a given session. These pooled manifolds therefore represent condition-averaged population geometry rather than simultaneously recorded single-trial dynamics. Statistical significance of pairwise trajectory relationships was assessed across recording sessions using Wilcoxon signed-rank tests, whereas uncertainty of effect-size estimates was quantified using bootstrap-derived confidence intervals (10,000 resamples). Resulting p-values were corrected using the Benjamini–Hochberg procedure. Robustness of the pooled geometry was further assessed using leave-one-session-out analyses combined with linear regression and Pearson correlation.

### Neural decoding analyses

Neural decoding analyses were performed using the Neural Decoding Toolbox (65) (NDT) implemented in MATLAB R2023b. Population vectors were constructed from spike counts of all available single-unit and multi-unit recordings pooled across recording sessions. Because neurons were not recorded simultaneously across sessions, decoding analyses were performed on pseudo-population datasets generated using the NDT framework.

Decoding was performed using a Poisson Naive Bayes classifier without additional feature preprocessing or firing-rate normalization. Classification performance was estimated using the cross-validator with 20 independent resample runs. The number of cross-validation folds was determined automatically for each analysis as the maximum feasible value supported by the minimum number of trials available across behavioral conditions, with a maximum of eight folds permitted. Decoder accuracy was computed independently for each temporal bin using spike counts measured in 100 ms windows advanced in 20 ms increments over the interval from −2 s to +1.5 s relative to behavioral onset. Statistical significance was assessed using 500 label-shuffled permutations to generate empirical null distributions of decoding accuracy, and p-values were estimated using the NDT framework. Decoder latency was defined as the first time point exhibiting significant decoding (p < 0.05) for at least five consecutive time bins. Decoding performance was evaluated using four behavioral classes (yawn, threat, bite, and drinking), yielding a theoretical chance level of 25%.

### Motor output mapping and M1-PM border identification via intracortical microstimulation

At the end of the recording sessions, intracortical microstimulation (ICMS) was applied at 542 of the 640 total recording sites, distributed across both hemispheres of the two monkeys, to verify the causal impact of local spiking activity on the motor output.

At each site, stimulation was delivered while the monkey was seated calmly in the primate chair with the head free, enabling to appreciate head, neck and axio-proximal movement in addition to the distal movements. Monopolar, biphasic trains of cathodic square-wave pulses were delivered via a constant-current stimulator (PlexStim, Plexon). Each train lasted 50 or 500 ms, with a single pulse width of 0.2 ms, a frequency of 200 Hz, and current intensities ranging from 7 μA to 150 μA. A site was attributed to the primary motor cortex if movements could be evoked with 50 ms ICMS trains at intensities of 25 μA or lower and if they were located within 2 mm of the presumed anatomical border between premotor and primary motor cortices, based on prior anatomo-functional studies (66).

Evoked movements were identified independently by two experimenters observing the animal during stimulation and were subsequently verified by offline analysis of video recordings synchronized to the ICMS pulses by simultaneously capturing the animal and an oscilloscope display that visualized the train of pulses. A site was assigned to a specific effector if stimulation at a given intensity consistently evoked the same movement, typically across at least three consecutive trials with identical parameters. Movements evoked by ICMS were categorized into the following effectors: *mouth/face* (facial movements including jaw, lip, tongue, periocular facial movements), *hand* (movements of fingers and/or wrist), *hand-mouth* (simultaneous movements of hand and mouth), and *head/axial* (movements of neck/head, trunk, shoulder or upper arm), *head/axial-mouth/face* (mouth/face movements occurring simultaneously with head/axial movements).

### Head-fixed yawn observation experiment

Head-fixed observation experiments were carried out by using selected videoclips depicting yawning or the same subjects before or after a yawn, to keep as much similar as possible the identity of the monkeys and the background between yawn video and the corresponding controls. Clips were analyzed using BORIS, annotating yawn onset, maximal mouth aperture, and yawn offset.

Neural activity was analyzed within a 3-s window starting 500 ms after clip onset for both yawn and matched control clips, thereby minimizing the contribution of transient visual responses associated with clip onset. Spike trains from each SU and/or MU were binned into 20-ms bins, smoothed with a Gaussian kernel (σ = 3), and averaged into five consecutive 500-ms epochs. The first epoch preceded yawn onset in the yawn condition and served as the baseline, whereas the remaining four epochs covered the period during which yawning occurred. The same temporal segmentation was applied to the matched control clips.

To determine whether a unit (single- or multiunit) was differentially modulated by the observation of yawning relative to matched control stimuli, firing rates were analyzed using a two-way repeated-measures ANOVA with Condition (Yawn, Control) and Epoch (five levels) as within-factors. A unit was classified as significantly modulated by yawn observation if at least one test epoch differed significantly from both baseline and the corresponding control epoch, with both comparisons indicating modulation in the same direction (i.e., facilitation or suppression).

## Acknowledgments

We thank F. Mastrandrea for drawings; A. Guida and L. Gioiosa for animal care and technical assistance; F. Biaggi and F. Leonardi for veterinary assistance and anesthesia; M. Pellegrini, R. Spanu and M. Delgrosso, for help in animal handling and data acquisition; L. Pedruzzi for providing macaque video recordings used to generate the experimental stimuli.

This work was supported by the European Research Council (ERC StG-2015 grant WIRELESS, 678307 to LB; ERC CoG-2020 grant EMACTIVE, 101002704 to LB; ERC PoC-2020 grant FUTURE-NHP, 957538 to LB), by the NEXTGENERATIONEU project MNESYS (PE0000006 - DN. 1553 11.10.2022) to LB, by the Italian Ministry of University and Research PRIN 2022 (22SP5K99) to MM, and by the Italian MUR grant CIRCEM R20NJ7BBA7 to LB.

