## Supplementary information for "Automatic orofacial behaviors remap within cortical substrates for voluntary motor control in freely moving primates"

**This PDF file includes:**

Figures S1 to S7
Tables S1 to S2
Legends for Movies S1 to S2

**Other supporting materials for this manuscript include the following:**

Movies S1 to S2

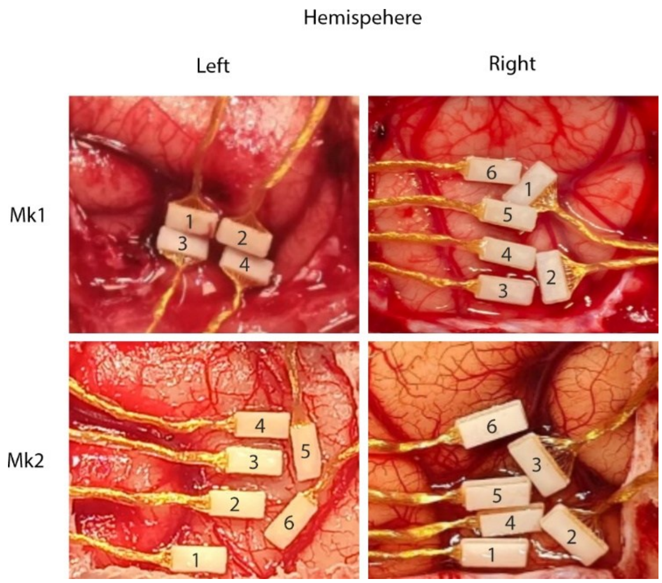


**Fig. S1. Implantation sites of Floating Microelectrode Arrays in Mk1 and Mk2.** Intraoperative photographs showing the left and right hemisphere of each monkey with the implanted arrays.

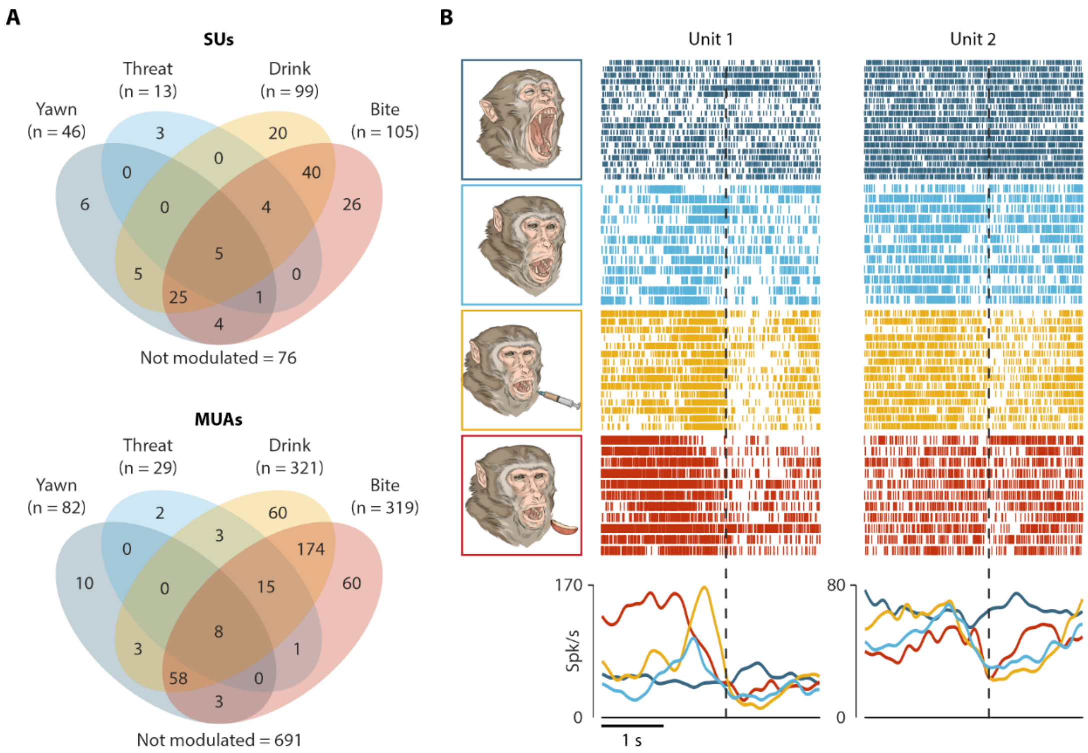


**Fig. S2. Units (single and multi) modulated during automatic and voluntary orofacial actions**. (A) Number of single units (top) and multiunits (bottom) units modulated during yawn, threat, drink, and bite. Non-modulated units are indicated below each diagram. (B) Example units modulated during threat displays that were often modulated during other voluntary orofacial behaviors, but not during yawning.

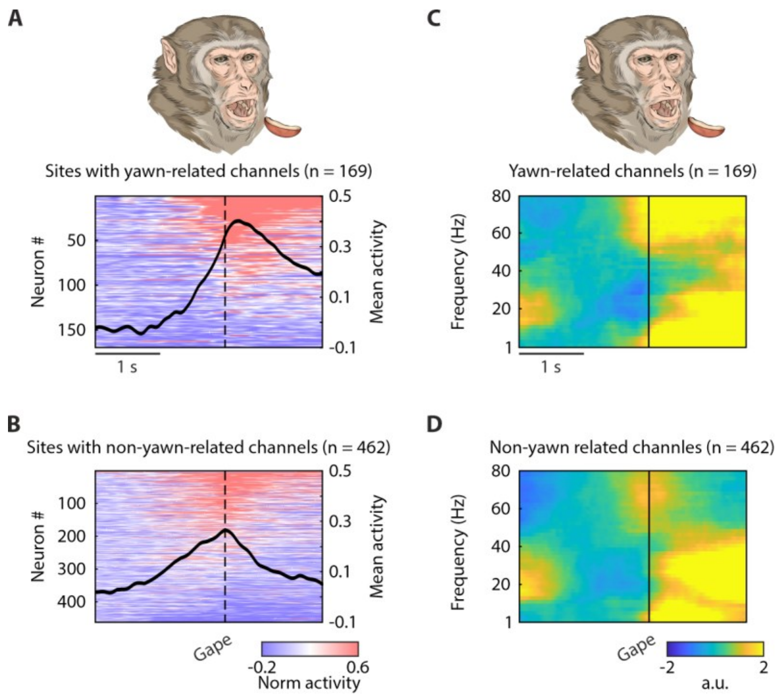


**Fig. S3. Yawn-related and non-yawn-related multiunit and LFPs activity during biting.** (A-B) Heatmaps and corresponding average time courses (black polylines) of normalized units for yawn-related (A) and non-yawn-related (B) units during biting. Vertical black lines indicate the alignment points, corresponding to the maximal mouth aperture. (C-D) Time course of LFP power modulation across the 1-200 Hz frequency range, averaged across sites with yawn-related (C) and non-yawn-related (D) units during biting.

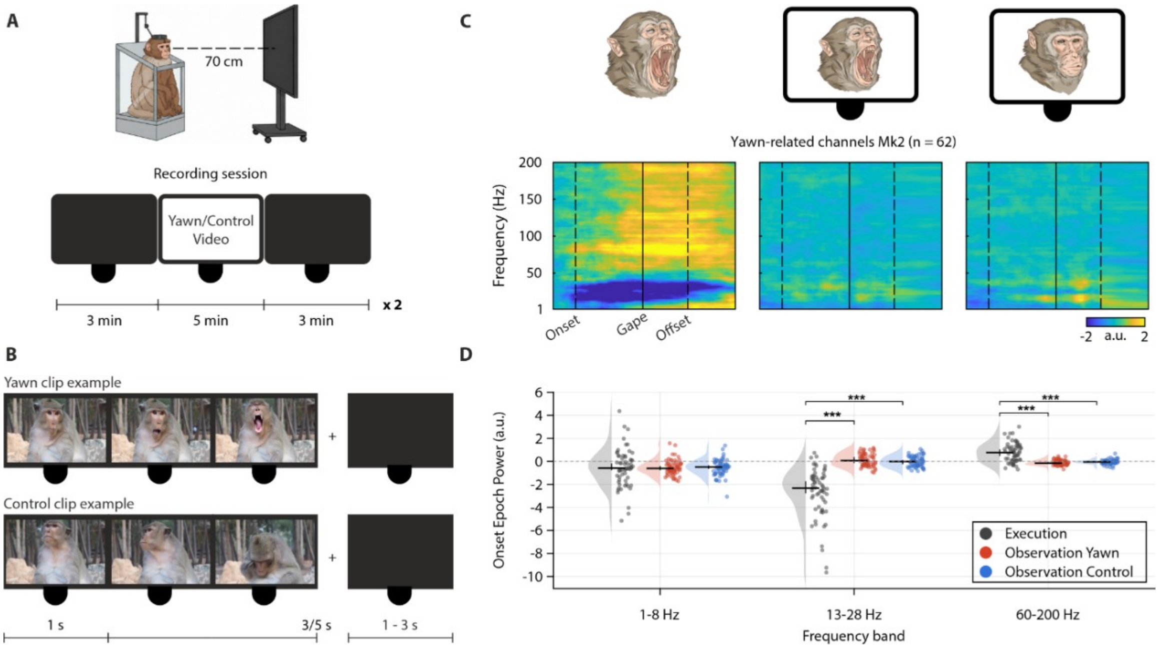


**Fig. S4. Head-fixed yawn observation paradigm.** (A) Top: experimental setup showing the monkey seated in the primate chair with the head fixed and the 32-inch monitor positioned 70 cm from the animal. Bottom: schematic representation of the recording session. Each session consisted of two experimental blocks (yawn and control, order randomized), each comprising a 3-min black screen, a 5-min video presentation, and a 3-min black screen. During the video presentation, the monkey viewed repeated presentations of video clips depicting familiar or unfamiliar conspecifics, separated by variable 1–3 s black-screen inter-trial interval, yielding a total stimulus duration of 5 minutes. (B) Timeline of an example yawn clip and its matched control clip. Each clip was followed by a black-screen inter-trial interval (1 - 3 s). In yawn clips, the conspecific remained visible for the first second after clip onset before initiating the yawn. Control clips were matched for subject identity, background and duration, but showed the same conspecific engaged in non-yawning behavior. We tested a total of 780 units (271 SUs and 509 MUAs) by applying a 2 x 5 repeated measures ANOVA (factor: Condition and Epoch) to each unit. Out of them, 24 units (3.1%; 7/271 SUs and 17/509 MUAs) exhibited a significant main effect of the factor epoch and a significant post-hoc effect between baseline and at least one of the tested epochs. Four units (0.5%; 0/271 SUs and 4/509 MUAs), none of which also exhibited a significant main effect of Epoch, showed a significant interaction effect with at least one of the test epochs, significantly and coherently (i.e. facilitated or suppressed) different from both baseline and the correspondent epoch of the control stimuli. Overall, this analysis revealed that only the 0.5% of the tested units exhibited a significant, but generally very weak, modulation during yawn relative to control stimuli, suggesting that no significant spiking output is generated during yawn observation from the explored motor and premotor cortical territory. (C) Population-averaged, baseline-normalized LFP spectrograms from yawn-related recording sites during yawn execution, observation of yawning, and observation of control videos. To ensure comparable sampling across conditions, analyses were restricted to matched recording sites from the same monkey, hemisphere, and arrays. Repeated trials, clips, and recording sessions were averaged within each physical recording site before population averaging. All spectrograms are displayed using identical frequency, time, and color scales (n = 62 matched recording sites per condition). (D) Site-level modulation of LFP power during the yawn onset epoch in the 1–8 Hz, 13–28 Hz, and 60–200 Hz bands. Each data point represents one physical recording site after averaging repeated measurements across trials, clips, and sessions. Horizontal bars indicate medians and vertical bars denote 95% bootstrap confidence intervals of the corresponding distribution. Overall differences among conditions were significant for the 13–28 Hz band (Kruskal–Wallis, $P = 8.31 \times 10^{-22}$) and 60–200 Hz band ($P = 9.72 \times 10^{-13}$), but not for the 1–8 Hz band ($P = 0.929$). Pairwise comparisons (paired Wilcoxon signed-rank tests with

Benjamini–Hochberg FDR correction) showed that differences were limited to the comparisons between yawn execution and observation conditions, with no significant effect in any frequency band for the comparisons between yawn and control stimuli. Statistical annotations indicate comparisons meeting both the FDR threshold ($q < 0.05$) and the predefined minimum effect-size criterion ($|\text{median difference}| \geq 0.2$ z-units).

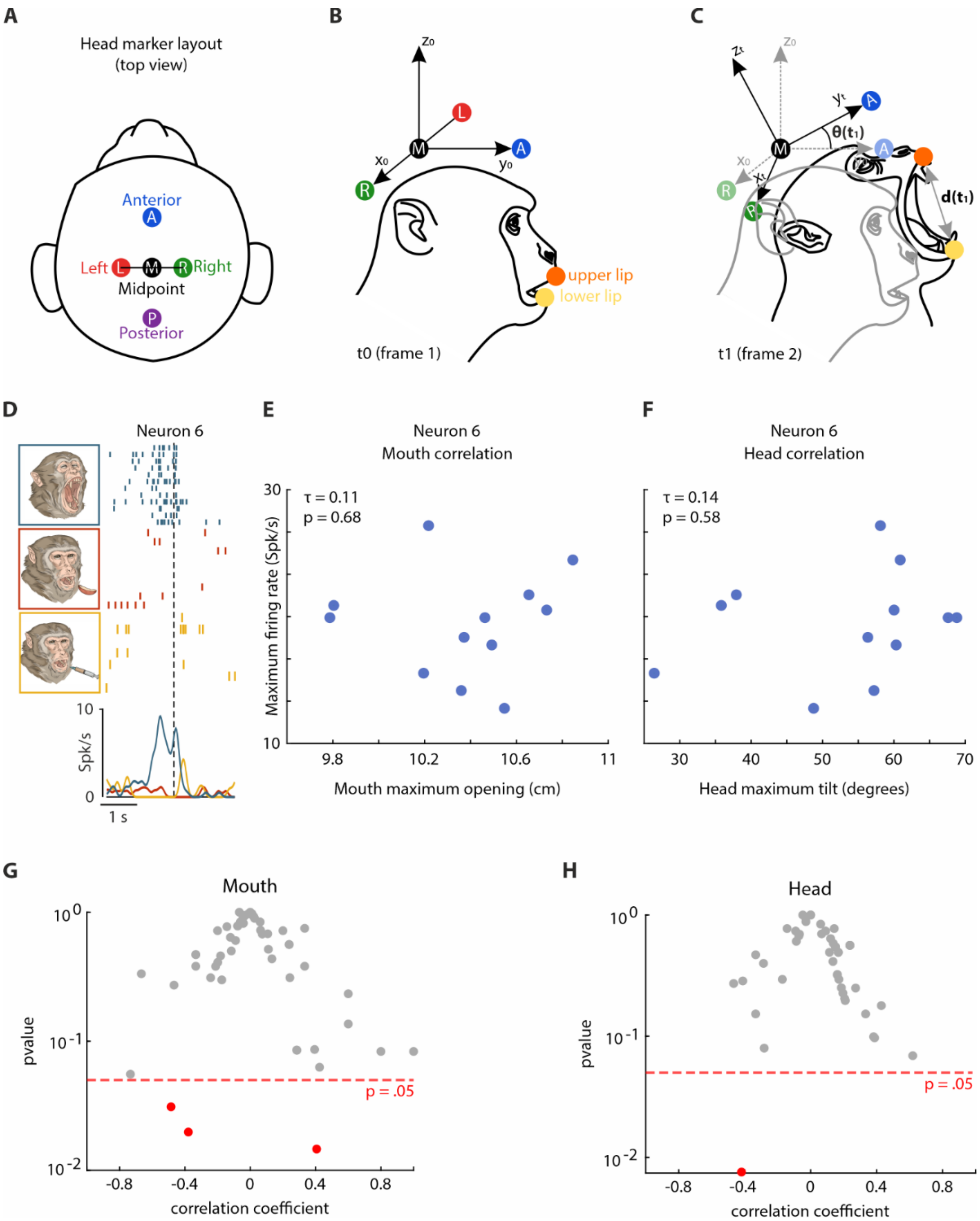


**Fig. S5. Kinematics analysis.** (A) Top view of the head marker layout mounted on the monkey headpost. (B) Head reference sagittal-plane frame at t0 (x0,y0,z0). The left-right axis (x0) is defined as the unit vector pointing from the left temporal marker position at t0, L(t0), to the right temporal marker position at t0 R(t0). The anterior-posterior axis (y0) is defined, depending on the marker configuration, as the unit vector pointing from the temporal midpoint at t0, M(t0), to the anterior marker at t0, or from the posterior marker at t0 to the temporal midpoint. The figure illustrates the former configuration (from the temporal midpoint to the anterior marker). The superior-inferior axis (z0) is defined as the unit vector obtained by normalizing the cross product of the left-right axis (x0 ) and the anterior-posterior axis (y0). The upper (orange dot), and lower (yellow dot) lips were marked at the midpoint of the upper and lower lips, respectively. (C) Head axis direction (y1) in the subsequent frame t1, was defined in the same manner as for t0: the left-right axis (x1) is defined as the unit vector pointing from the left temporal marker position at t1, L(t1), to the right temporal marker position at t1 R(t1). The anterior-posterior axis (y1) is defined as the unit vector pointing from the temporal midpoint at t1, M(t1), to the anterior marker at t1, the superior-inferior axis (z1) is defined as the unit vector obtained by normalizing the cross product of the left-right axis (x1) and the anterior-posterior axis (y1). To isolate upward-downward head motion (pitch), the component of the current head axis (y1) aligned with the fixed baseline left-right axis (x0) was subtracted, thereby projecting the head axis onto the baseline sagittal plane. The instantaneous head tilt angle is computed within the baseline sagittal plane as the angle between the projected head axis (y1) and the initial head direction ($y_0$). Mouth opening is defined as the Euclidean distance between the upper and lower lip markers ($d(t1) = ||u(t1) - l(t1)||$). (D) Raster plot and peri-event time histogram of an example yawn-related neuron. Peak neuronal activity coincides with the alignment point, corresponding to the time of maximum mouth opening and maximum head elevation. (E-F) Scatter plots showing the relationship between the peak of the firing rate (spikes/s) and maximum mouth opening (E) or maximum head elevation (F) across yawn trials (blue dots) for the example neuron shown in D. (G–H) Summary plots of the Kendall's rank correlation analysis between neuronal response and mouth (G) and head (H) kinematics. Each dot represents one neuron and is plotted according to its Kendall's correlation coefficient ($\tau$) and associated p-value. Neurons showing a significant correlation with maximum mouth opening ($n = 4$) or maximum head elevation ($n = 1$) ($p < 0.05$) are highlighted in red, whereas non-significant neurons are shown in gray. The dashed red horizontal line indicates the significance threshold ($p = 0.05$).

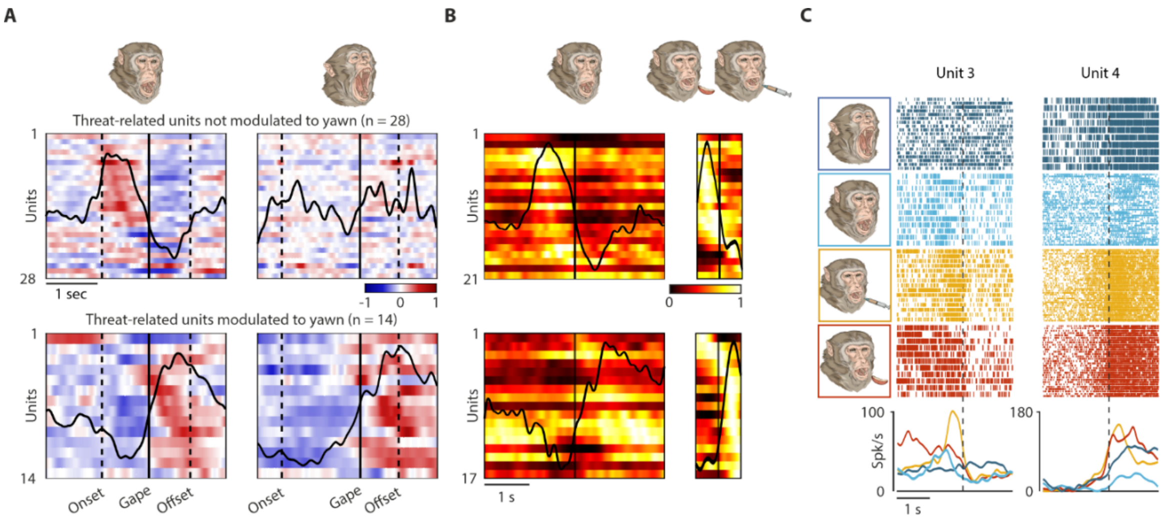


**Fig. S6. Threat-related cortical activity and its relationship with yawning and voluntary orofacial actions.** (A) Heatmaps and corresponding average time course (black polyline) of the net-normalized activity of threat-related units during threat and yawn events, shown from -2 to 1.5 relative to mouth gape (black vertical line). Conventions as in Fig. 2E. (B) Net-normalized activity of threat-related units peaking during the mouth-opening (top; n = 21) and mouth-closing (bottom; n = 17) phase of threat, shown from -2 to 2 s during threat and from -0.5 to 0.5 during voluntary mouth action. Conventions as in Fig. 3A, C. (C) Examples of a threat-related units unmodulated (Unit 3) and modulated (Unit 4) to yawning, showing a synergistic modulation with voluntary orofacial actions (i.e. bite and drink) during mouth opening (Unit 3) and mouth closure (Unit 4).

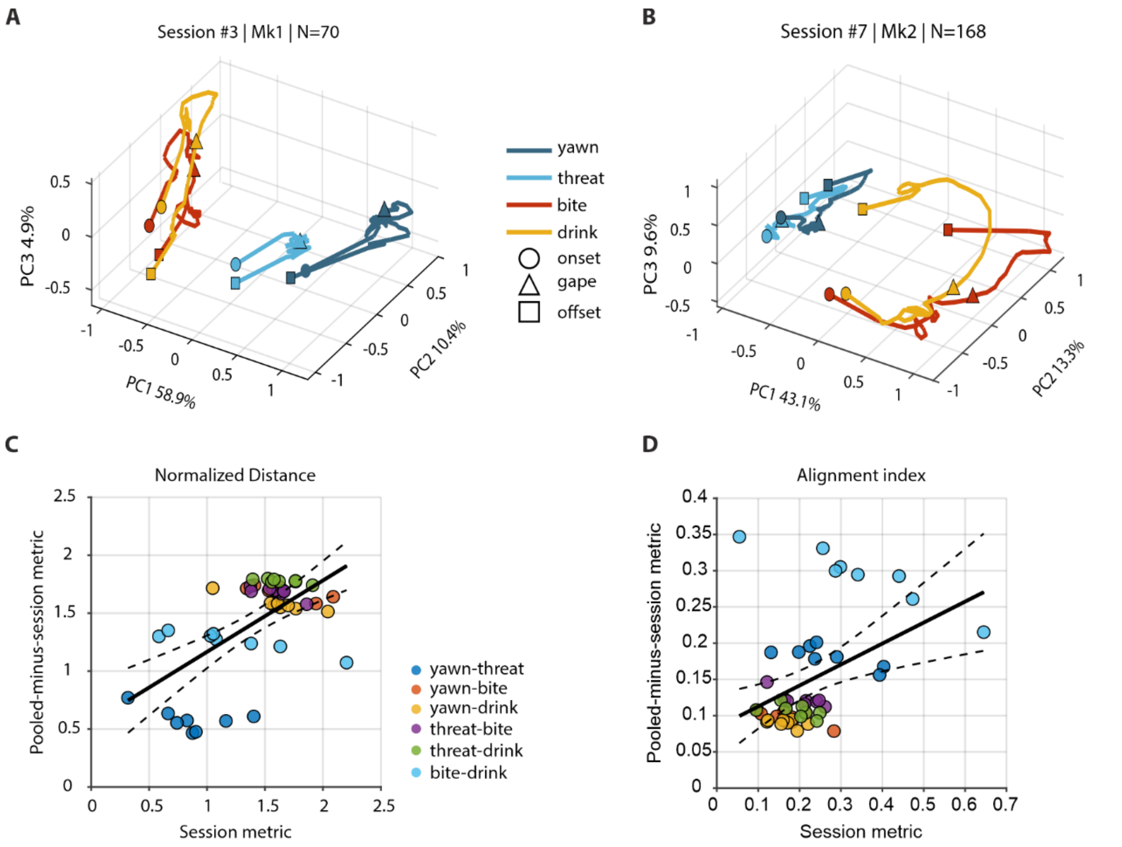


**Fig. S7. Session-level validation of latent population geometry and robustness of pooled pseudo-population trajectories.** (A–B) Representative three-dimensional latent trajectories reconstructed independently from simultaneously recorded neurons within two example recording sessions (Session #3, Mk1, N = 70 units; Session #7, Mk2, N = 168 units). Population activity from all available units (SUA+MUA) was projected onto the first three principal components of the phase-normalized neural manifold. Trajectories depict the temporal evolution of neural population activity during yawning (blue), threat (red), biting (green), and drinking (purple), from action onset (circles), through the phase-normalized maximal-effector state (triangles), to action termination (squares). (C) Leave-one-session-out (LOSO) validation of normalized trajectory distance across all action pairs. For each session, pairwise trajectory distances were computed using only neurons simultaneously recorded within that session and compared against the corresponding metric obtained from the pooled dataset after excluding that session. The significant positive relationship between session-specific and pooled-minus-session values (linear regression: $R^2 = 0.37$, $p = 4.01 \times 10^{-6}$; Pearson $r = 0.61$, $p = 4.01 \times 10^{-6}$) demonstrates that pairwise trajectory geometry is reproducible across recording sessions and is not driven by any individual dataset. Across-session analyses identified the yawn–threat pair as exhibiting the smallest trajectory distance (mean = 0.86, $p = 3.91 \times 10^{-3}$, FDR-corrected $q = 0.023$), indicating that yawning and threat followed the most similar population dynamics among all behavioral pairs. (D) LOSO validation of the alignment index, which quantifies overlap between action-specific latent subspaces. Session-derived alignment values significantly predicted alignment values obtained from the pooled dataset excluding that session (linear regression: $R^2 = 0.18$, $p = 3.07 \times 10^{-3}$; Pearson $r = 0.42$, $p = 3.07 \times 10^{-3}$). Significant relationships in both panels C and D indicate that the latent geometry underlying yawning, threat, biting, and drinking is consistently observed across independent recording sessions. Across sessions, bite–drink exhibited the highest subspace alignment (mean = 0.35, $p = 0.0195$), followed by yawn–threat (mean = 0.27, $p = 0.0273$), consistent with partially shared latent representations between these behavioral pairs.

| Monkey | Hemisphere | Arrays | Sessions per 32 ch array | Channels sampled |
|---|---|---|---|---|
| **Mk1** | Left | 1, 2, 3, 4 | 4 | 512 |
| | Right | 1, 2 | 1 | 64 |
| | Right | 3, 4 | 3 | 192 |
| | Right | 5, 6 | 1 | 64 |
| **Mk2** | Left | 2, 3, 4, 5 | 1 | 128 |
| | Right | 1, 3 | 4 | 256 |
| | Right | 2, 4 | 3 | 192 |
| | Right | 5, 6 | 1 | 64 |
| **Total** | | | | **1472** |

**Table S1. Detailed information regarding monkeys, hemispheres and arrays across all the recording sessions.** For each 32-channel array (see Figure S1), the number of sessions in which it was sampled and the corresponding total number of channels sampled (or resampled) is reported. From half of the sites (n = 320) we recorded only once, whereas the remaining 320 sites were sampled across 3-4 distinct sessions.

| Actions | Definition of the scored event |
|---|---|
| 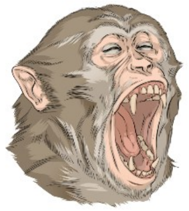 | **Yawn**<br>The monkey opens its mouth, takes a deep inspiration followed by a slow expiration, and then closes its mouth. The marked event corresponded to the moment of maximal mouth opening before mouth closure began. Action onset (i.e. the beginning of mouth opening) and offset (i.e. the completion of mouth closure) were also marked. |
| 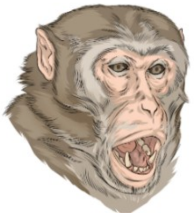 | **Threat**<br>The monkey opens its mouth, with or without exposing its canines, while staring at the experimenter in response to the experimenter's mouth-open threat gesture. The marked event corresponded to the moment of maximal mouth opening before mouth closure began. Behavioral onset (i.e. the beginning of mouth opening) and offset (i.e. the completion of mouth closure) were marked. |
| 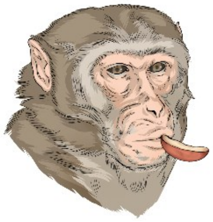 | **Bite**<br>The monkey places food into its mouth either by closing its lips around food delivered directly by the experimenter using a stick or, when this behavior was not available, by bringing food to the mouth with the contralateral hand after grasping it from the floor or from a thread. The marked event corresponded to the moment when the food was fully taken into the mouth (i.e., lip closure around the food in experimenter-delivered trials, or complete placement of the food into the mouth in self-delivered trials). Behavioral onset (i.e. the moment when the mouth began to open in preparation for food intake), offset (i.e. the moment when the mouth reopened or the arm moved away after food placement) and mouth maximum opening (i.e. the moment of maximal mouth aperture while the monkey approached the experimenter's stick or brought the food toward the mouth) were also marked. |
| 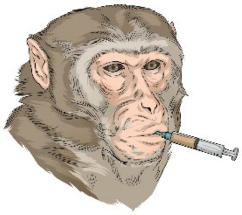 | **Drink**<br>The monkey drinks juice by opening its mouth and closing its lips around the tip of a plastic syringe presented directly by the experimenter. The marked event corresponded to the moment when the lips closed around the syringe spout. Behavioral onset (i.e. the moment when the mouth began to open as the monkey approached the syringe), offset (i.e. the moment when the mouth opened again as the monkey detached from the syringe) and mouth maximum opening (i.e. the moment of maximal mouth aperture while the monkey approached the syringe) were marked. |

**Table S2. Ethogram of the investigated orofacial actions.**

**Movie S1 (separate file).** Example of lips and head tacking during yawn.
**Movie S2 (separate file).** Example of latent trajectories evolving through a three-dimensional state space.
